# Structural basis for dimerization, catalytic regulation, and substrate selectivity in S9D proteases

**DOI:** 10.64898/2026.01.21.700880

**Authors:** Jacqueline J. Ehrlich, Pratyush Routray, Louis Enns, Klaas J. van Wijk, Toshimitsu Kawate

**Author notes:** Corresponding authors: Klaas J. van Wijk, School of Integrative Plant Science, Plant Biology Section, Cornell University, 332 Emerson Hall, Ithaca, NY 14853; Toshimitsu Kawate, Department of Molecular Medicine, Cornell University, C4-151 VMC, 930 Campus Rd, Ithaca, NY 14853, Email: tosh.

## Abstract

S9 proteases are widely distributed across the tree-of-life and play essential roles in protein processing. However, the structural and mechanistic basis for protease activity in the S9D subfamily has remained unknown. Here, we report the first high-resolution cryo-EM structures of an S9D protease, chloroplast glutamyl endopeptidase (CGEP) from *Arabidopsis thaliana*, expressed in plants and bacteria. CGEP adopts a dimeric architecture stabilized by two distinct interfaces: hydrophobic interactions between catalytic domains and an interdomain β-sheet linking the cap and catalytic domains. These interactions create a rigid scaffold that supports a hinge loop, which acts as a steric gate to restrict substrate access and confine catalytic activity to the closed conformation. Unlike S9A-B-C proteases, CGEP maintains an intact catalytic triad in both open and closed states, relying on hinge-loop gating rather than catalytic disruption for regulation. Structural analysis and mutagenesis reveal that the hinge loop forms a conserved pocket favoring glutamate side chains, explaining CGEP’s strong glutamate preference at cleavage sites. Together, these findings uncover a unique regulatory paradigm for S9D proteases and provide a structural framework for understanding substrate selectivity and dimerization.

**Significance statement:** This study provides the first structural and mechanistic insights into plant S9D proteases, revealing a unique regulatory paradigm that combines hinge-loop gating with substrate-specific recognition. By uncovering how CGEP maintains catalytic integrity while restricting activity to the closed conformation and explaining its strong glutamate preference, these findings advance our understanding of protease diversity and open new avenues for engineering proteases with tailored specificity for agricultural and biotechnological applications.

## Introduction

The S9 family of serine proteases comprises a diverse group of serine-dependent peptidases classified into four subfamilies: S9A, S9B, S9C, and S9D (Kiss-Szeman et al. 2019; Rawlings et al. 2018; Rawlings and Bateman 2021). Their catalytic triad residues exist in the order Ser-Asp-His within the primary sequence. Each subfamily is characterized by a distinct motif surrounding the active-site serine: GGSXGGLL (S9A), GWSYGGY (S9B), GGSYGG (S9C), and GGHSYGAFMT (S9D), where X is typically Asn or Ala (Rawlings et al. 2018). S9 proteases exhibit diverse cleavage specificities, often targeting peptide bonds C-terminal to residues such as Pro, Arg, Lys, Glu or acetylated amino acids. These enzymes primarily cleave short oligopeptides (<30 amino acids, about 3.3 kDa) rather than larger proteins (Kiss-Szeman et al. 2019). Members of subfamilies S9A–S9C are known by various names depending on species and activity, including prolyl oligopeptidase (POP), prolyl endopeptidase (PEP), dipeptidyl peptidase IV (DPP4), and acyl-aminoacyl peptidase (AAP) (Polgar 2002). In *Arabidopsis thaliana* and other higher plants, representatives of S9A (AARE1 and AARE2; AT4G14570 and AT5G36210) and S9B (AT5G24260) have been identified, whereas no S9C homologs are present (Hoernstein et al. 2023; Nakai et al. 2012; Yamauchi et al. 2003).

Tertiary structures have been solved for several members of the S9A subfamily (*e.g.* PDB IDs: 1QFM (Fulop et al. 1998), 6CAN and 5T88 (Ellis-Guardiola et al. 2019) 9HJI and 9HJJ (Batra et al. 2025)), the S9B subfamily (*e.g.* PDB IDs: 2ECF (Nakajima et al. 2008), 1J2E (Hiramatsu et al. 2003)), and the S9C subfamily (*e.g.* PDB IDs: 7EP9 (Wang et al. 2021), 5YZM, 5YZN, 5YZO, 6IGP, 6IGR, and 6IGQ (Yadav et al. 2019)). In contrast, no structure is currently available for members of the S9D subfamily. All S9A-C proteins with known structures share a conserved architecture comprising an N-terminal seven- or eight-bladed β-propeller, formed by four-stranded antiparallel β-sheets, and a C-terminal α/β/α sandwich peptidase domain. Interestingly, S9 proteases exhibit diverse oligomeric states even within the same subfamily, with observed structures forming as monomers, dimers, tetramers, or hexamers (Table S1). The β-propeller domain of certain S9 proteases regulates substrate access to the catalytic site in the C-terminal α/β/α sandwich peptidase domain (Batra et al. 2025; Harmat et al. 2011; Kiss-Szeman et al. 2019). Conformational changes in this domain can also influence the enzyme’s activation state, as opening of the β-propeller domain appears to disrupt the catalytic triad in certain species (Harmat et al. 2011; Kiss-Szeman et al. 2019). Differences in oligomerization states among subfamilies (S9A monomer, S9B tetramer, S9C dimer) are thought to arise from a “sticky” region within the repeat β-sheet of the hydrolase domain, often called “sticky β-edge”, where this terminal β-strand mediates oligomerization (Kiss-Szeman et al. 2019).

Phylogenetic analyses reveal that S9D proteases are present not only in higher plants (both monocots and dicots), but also in early land plants such as mosses and lycopods, in red and green algae, in cyanobacteria, and in Gram-negative non-photosynthetic prokaryotes, including α-, β-, and γ-proteobacteria as well as flavobacteria (Bhuiyan et al. 2020). In contrast, S9D members appear to be absent from other major branches of the tree-of-life (*e.g.* archaea, fungi, animals). In a recent study, we characterized the proteolytic cleavage specificity of a chloroplast-localized S9D member in the plant *Arabidopsis thaliana* (AT2G47390) that we named Chloroplast Glutamyl Peptidase (CGEP) (Bhuiyan et al. 2020). Using large peptide libraries and mass spectrometry, we demonstrated that recombinant CGEP exhibits strict specificity for cleavage on the C-terminal side of glutamate, regardless of the identity of adjacent residues. This precise substrate preference aligns with earlier biochemical observations from chloroplast extract assays, where glutamyl aminopeptidase activity was detected but the responsible protease remained unidentified (Forsberg et al. 2005; Laing and Christeller 1997). Recombinant CGEP partially degrades 25 kD β-casein but completely degrades 10 kD insulin and smaller peptides through both exo- and endopeptidase cleavage. Interestingly, *Arabidopsis* CGEP undergoes complete autocatalytic C-terminal cleavage at E946, both *in vivo* and *in vitro (*recombinantly expressed in *E. coli)*, effectively removing the last 15 residues from its C-terminus. A conserved motif (A[S/T]GGG[N/G]PE946) located immediately downstream of E946 was identified in dicotyledonous species, but this region was altered in monocotyledonous species (ASGG[S/G]A[P/A][C/R]E). *In vivo* complementation in a *cgep clpr2-1* Arabidopsis background using either a catalytically inactive variant (CGEP-S781R) or a catalytically active variant lacking C-terminal processing (CGEP-E946A-E949A-E951A) demonstrated the physiological importance of both CGEP peptidase activity and its autocatalytic cleavage. CGEP homologs in photosynthetic and non-photosynthetic bacteria lack this C-terminal prosequence, suggesting it is a recent functional adaptation in plants. Structural modeling, using the best available structural template at the time (S9B dipeptidyl aminopeptidase IV - PDB:2ECF (Nakajima et al. 2008)), indicated that C-terminal processing may increase the upper substrate size limit by improving access to catalytic cavity (Bhuiyan et al. 2020). Although these studies revealed important characteristics of CGEP, the underlying mechanisms remain unresolved due to the absence of a high-resolution structure for the S9D subfamily. Homology models using the existing S9A-C proteins are not reliable because of the low sequence similarity—only 11-13% identical to other *Arabidopsis thaliana* S9 proteases (Fig. S1). High-resolution structural information is critical because it enables precise visualization of the catalytic site, substrate-entry sites, and dynamic regions that govern enzyme specificity and regulation.

The current study presents the first high-resolution structures of an S9D protease, determined by cryogenic electron microscopy (cryo-EM) using both plant-purified CGEP and recombinant CGEP expressed in *E. coli*. We identified both conserved and unique features of the S9D subfamily. Furthermore, targeted mutagenesis guided by these structural insights revealed key determinants of subtype-specific substrate recognition, shedding light on the molecular mechanisms underlying S9D function. These findings provide a structural framework for understanding how S9D proteases contribute to chloroplast protein homeostasis and may inform strategies to manipulate proteolytic activity for improving plant stress resilience and metabolic efficiency.

## Results

### CGEP assembles into a dimer featuring side openings for substrate access

To gain structural insights into the under-characterized S9D subfamily, we performed cryo-EM single-particle reconstruction of CGEP. We overexpressed C-terminal StrepII-tagged CGEP in *Arabidopsis thaliana* to preserve its native folding environment in the chloroplast. Based on our previous studies indicating that the last 15 amino acids of the C-terminal domain (CTD) (Fig. 1A) undergo autocleavage, we utilized transgenic *Arabidopsis* expressing CGEP with a mutation at the nucleophilic serine residue (S781R) within the catalytic triad to prevent cleavage of the CTD. Size-exclusion chromatography (SEC) analysis showed that the StrepII-affinity purified protein sample was monodisperse and mostly dimeric (Fig. S2A and B), although a fraction of CGEP appeared to form a monomer. We note that endogenous Arabidopsis CGEP is also a dimer (Bhuiyan et al. 2020). Using this protein sample purified from plant tissues, we obtained cryo-EM maps of dimeric CGEP at global resolution ranging from 3.1 to 3.6 Å, sufficient for reliable atomic model building for residues 110 and 915 (residues 696-705 are missing in the open conformation) (Fig. S3-5, Table S2).

**Figure 1.**
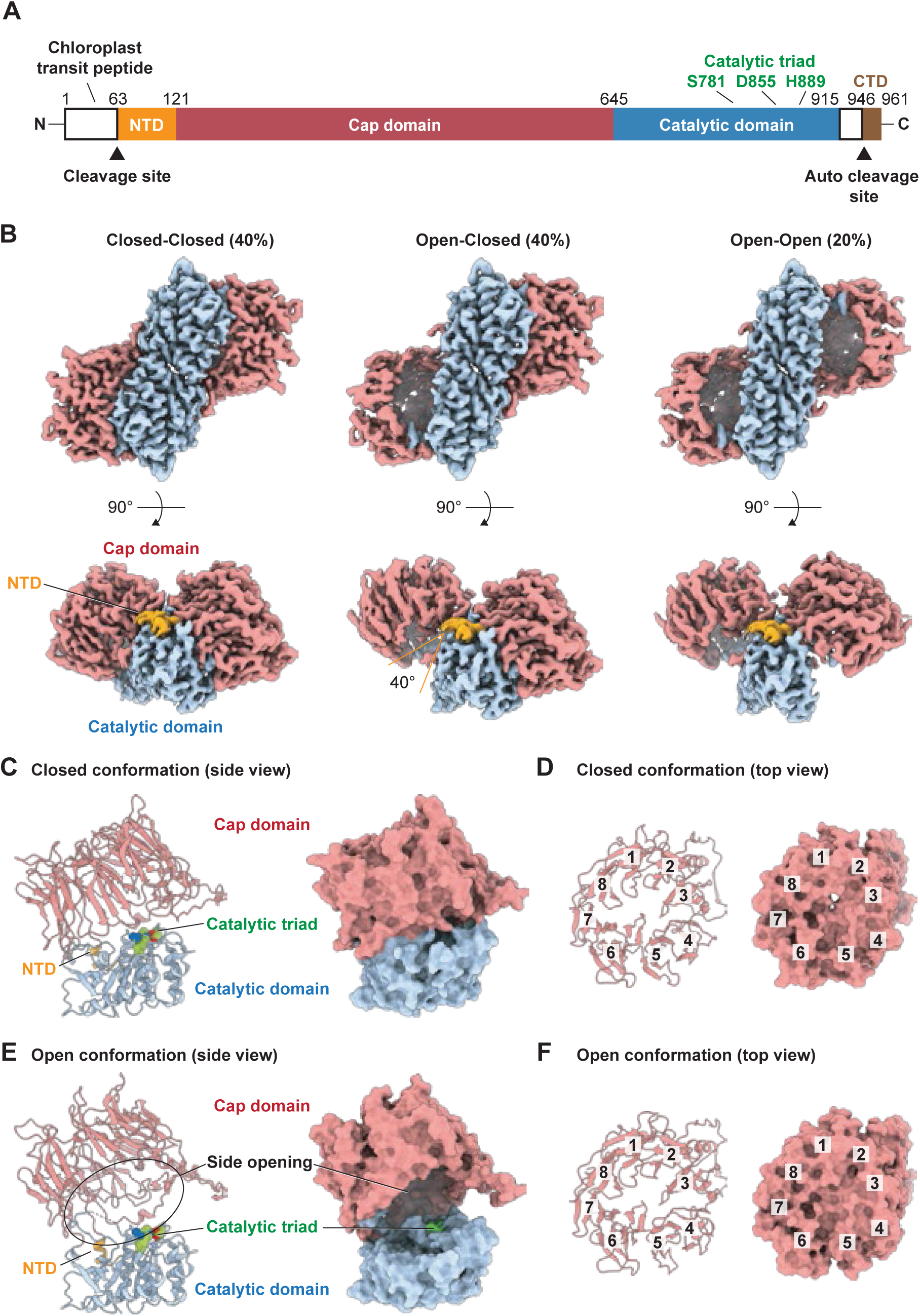
Dimeric CGEP reveals a side opening for substrate entry. (A) Schematic diagram of CGEP domains, highlighting the cleaved N-terminal chloroplast transit peptide, the N-terminal domain (NTD), cap domain, catalytic domain, and C-terminal domain (CTD) with its autocatalytic cleavage site (E946). The catalytic triad is shown in green. (B) Cryo-EM maps of CGEP S781R purified from transgenic *Arabidopsis* in three distinct configurations, with each domain colored as the diagram above. (C and D) Cartoon (left) and surface (right) representations of the closed conformation in side and top views. (E and F) Cartoon (left) and surface (right) representations of the open conformation inside (left) and top (right) views. Individual β-propellers are numbered in the top views (D and F).

Dimeric CGEP forms three distinct states: closed–closed (40%), closed–open (40%), and open–open (20%) (Fig. 1B). The major distinction among the dimeric conformations lay in the openness of the cap domain, while the catalytic domains were fully superimposable (RMSD = 0.64 Å). This observation suggests that substrate access in CGEP is primarily regulated by cap domain opening, whereas dimer formation does not play a direct role in this process.

The cap domain of CGEP adopts an eight-bladed β-propeller architecture, similar to that observed in S9B proteases (Fig. 1C-F, S6) (Kiss-Szeman et al. 2019). Consistent with other S9 subfamilies, the N⍰terminal domain (NTD) contributed to the architecture of the α/β/α sandwich catalytic domain (Fig. 1C, S6). Each lobe of the dimeric CGEP exhibits either a closed or open conformation, with the cap domain hinging away from the catalytic domain by approximately 40° in the open state (Fig. 1B-C and E). This is notable because dynamic cap movements are typically associated with monomeric S9 proteases (Kiss-Szeman et al. 2019). In the closed conformation, the cap and catalytic domains pack tightly against each other, leaving minimal space between them (Fig. 1C). The β⍰propeller core is similarly packed in both the open and closed states (Fig. 1D and F), effectively limiting direct access to the active site. These observations suggest that CGEP substrates are likely to reach the catalytic triad through side openings (Fig. 1E).

### A spacious cap-domain cavity facilitates processing of large substrates

CGEP exhibits partial degradation of the 25⍰kDa β⍰casein yet fully hydrolyzes 10⍰kDa insulin and smaller peptides through both exo11 and endopeptidase cleavage mechanisms (Bhuiyan et al. 2020). The presence of wide lateral entry openings (∼40Å x∼60Å) is consistent with this range of substrate sizes. In addition, CGEP features a spacious cap-domain cavity capable of accommodating a sphere with a 17 Å radius in both its closed and open conformations (Fig. 2A and B), implying that a globular substrate up to ∼10 kDa can be fully enclosed even when the enzyme is in the closed state. This architectural feature facilitates the digestion of relatively long peptides or small folded proteins such as insulin. Notably, the cross⍰sectional area of the cap⍰domain cavity is comparable to that of the S9B family protease DPP-IV, which is capable of cleaving full⍰length proteins of up to 10 kDa, including chemokines, CSFs, and interleukins (Ou et al. 2013; Zhu et al. 2003) (Fig. S7). Interestingly, conspicuous cryo⍰EM densities were observed only in the closed conformation, potentially representing substrate molecules captured during digestion (Fig. 2).

**Figure 2.**
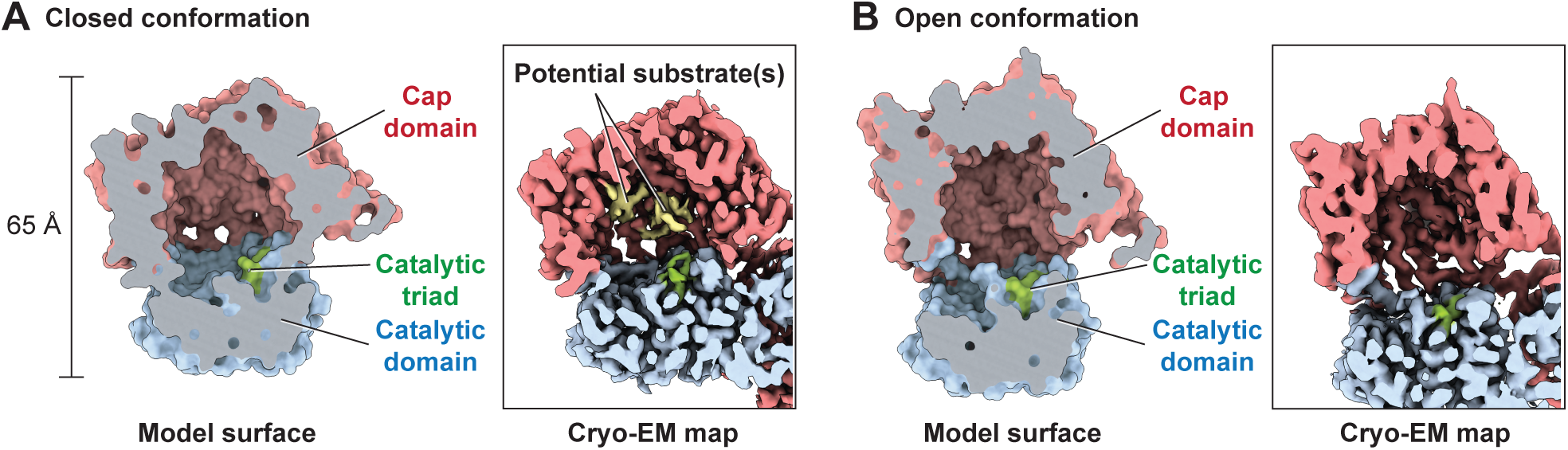
A spacious cap domain accommodates a large substrate. (A and B) Sagittal sections of CGEP S781R in the closed (A) and open (B) states. Model surface (left) and cryo-EM map (right) are shown. Each domain and the catalytic triad are colored as in Figure 1. Potential substrates (yellow) are observed only in the closed conformation. Each domain colored as in the diagram shown in Fig. 1A.

### Dimerization loop in the cap domain holds the dimer together

While dimerization is common among S9B proteases and some S9C proteases (Table S1), CGEP employs a notably distinct mode of dimer assembly. We identified two major dimer interfaces in the CGEP structures. The first interface is formed between neighboring catalytic domains (Fig. 3A and B) and is stabilized through hydrophobic interactions involving the side chains of W825, V832, and F837. This interface remains unchanged during cap opening and closing. The second dimer interface involves a loop in the cap domain spanning residues I276–D315, hereafter referred to as the “dimerization loop.” Although this loop undergoes substantial conformational rearrangement during cap opening, residues K294–N298 consistently form an antiparallel β⍰sheet with the eighth β⍰strand in the catalytic domain (Fig. 3C and D). This unique cross⍰domain interaction seamlessly links the cap domain of one subunit to the catalytic domain of the other. The eighth β⍰strand in the catalytic domain corresponds to the well⍰known “sticky β⍰edge,” which plays a central role in oligomerization among S9 proteases (Kiss-Szeman et al. 2019). In CGEP, this sticky β⍰edge is shielded by the dimerization loop in both the closed and open conformations, preventing higher⍰order oligomerization or aggregation.

**Figure 3.**
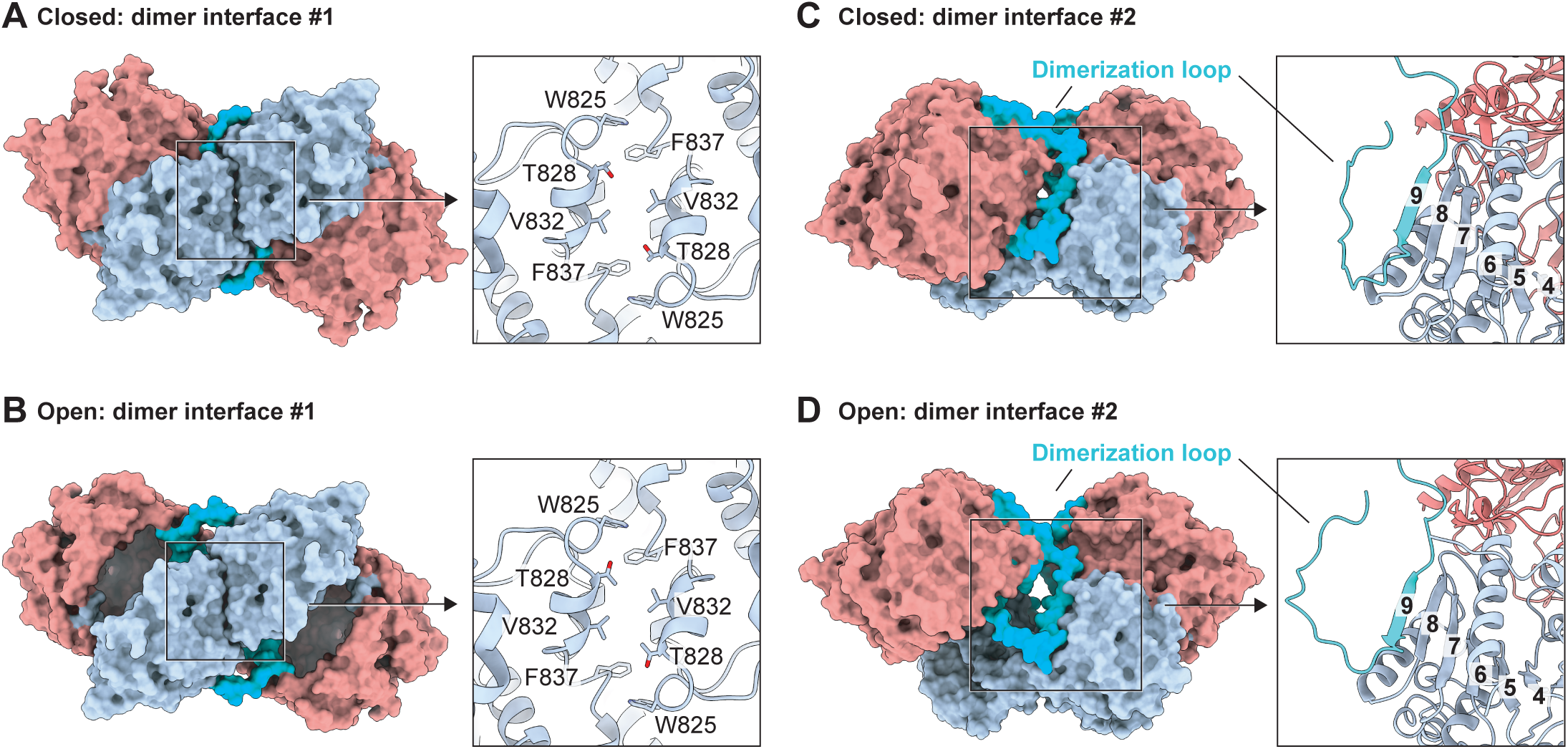
Two distinctive dimer interfaces. (A and B) Dimer interface mediated by hydrophobic interactions between neighboring catalytic domains in the closed (A) and open (B) states. (C and D) Dimer interface mediated by β-sheet formation between the cap and catalytic domains of different subunits in the closed (C) and open (D) states. Numbers indicate β-strands, with strand eight representing the sticky β-edge and strand nine representing the residues in the dimerization loop (cerulean). Left: surface view of CGEP S781R structure; Right: cartoon and stick representation of the boxed area. Each domain colored as in the diagram shown in Fig. 1A.

### Hinge loop between cap and catalytic domains functions as a steric gate for substrate access

To investigate how CGEP regulates its catalytic activity, we first compared the catalytic triad in the closed and open conformations. The positions of S781, D855, and H889 are essentially superimposable, indicating that the catalytic triad remains fully intact in both states (Fig. 4A, B). This behavior contrasts with other S9 proteases that undergo catalytic triad distortions during cap⍰opening motions. For example, in *Pyrococcus furiosus* POP, the catalytic histidine shifts by approximately 10 Å upon cap opening, disrupting the alignment of the triad and rendering the enzyme inactive in that conformation (Ellis-Guardiola et al. 2019). These observations suggest that CGEP employs a distinct mechanism to control its catalytic activity.

**Figure 4.**
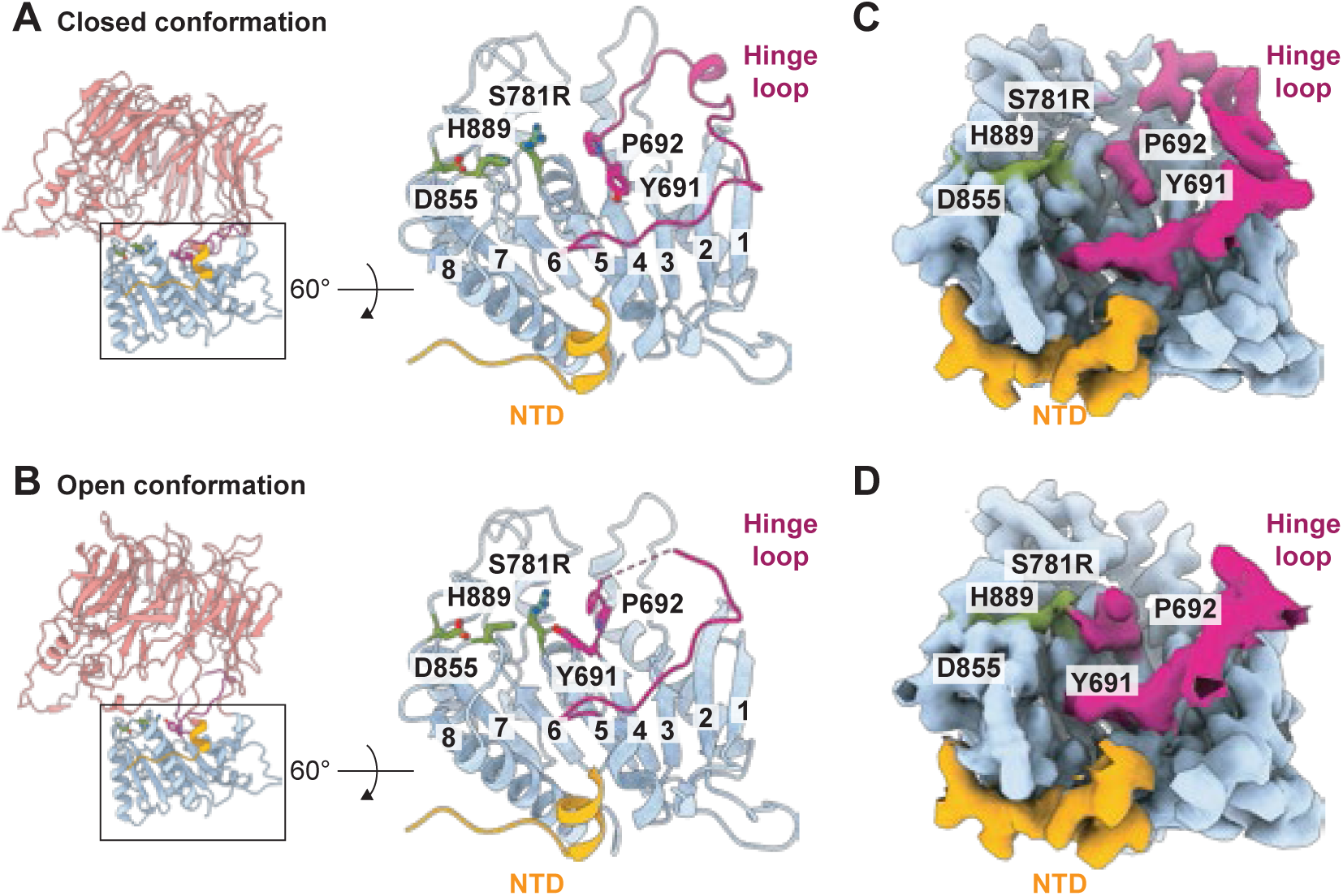
Comparison of the catalytic triad and hinge loop between closed and open states. (A and B) Cartoon representation of the CGEP S781R catalytic domain viewed from the top. Designated residues are shown in stick representation, and β-strands are numbered, with strand eight representing the sticky β-edge. Closed (A) and open (B) conformations are shown. (C and D) Cryo-EM maps of the catalytic domain viewed from the same angle as in (A) and (B). Closed (C) and open (D) conformations are shown.

We next compared the surrounding area and found that the loop in the catalytic domain that bridges the cap domain, spanning residues W689–S718 and referred to here as the “hinge loop,” undergoes a substantial conformational rearrangement during cap opening (Fig. 4A–D). Notably, residues Y691 and P692 flip toward the catalytic triad in the open conformation, likely obstructing substrate access to S781. This implies that substrate digestion occurs predominantly in the closed conformation, consistent with other S9 proteases that feature a mobile cap domain (Kiss-Szeman et al. 2019).

Because the S781R mutation might influence the hinge⍰loop conformation in the open state, we also recombinantly expressed and purified N-terminal His6-tagged wild⍰type *Arabidopsis* CGEP (N-terminal chloroplast signal peptide (cTP) was removed) from *E. coli* and performed cryo⍰EM single⍰particle reconstruction (Fig. S8-11 and Table S3). Consistent with the S781R structure, this wild⍰type enzyme displayed a superposable catalytic triad and showed restricted substrate accessibility in the open conformation due to hinge⍰loop rearrangement (Fig. 5A–D). These results confirm that the hinge loop plays a central role in limiting substrate entry to the catalytic triad when the cap is open.

**Figure 5.**
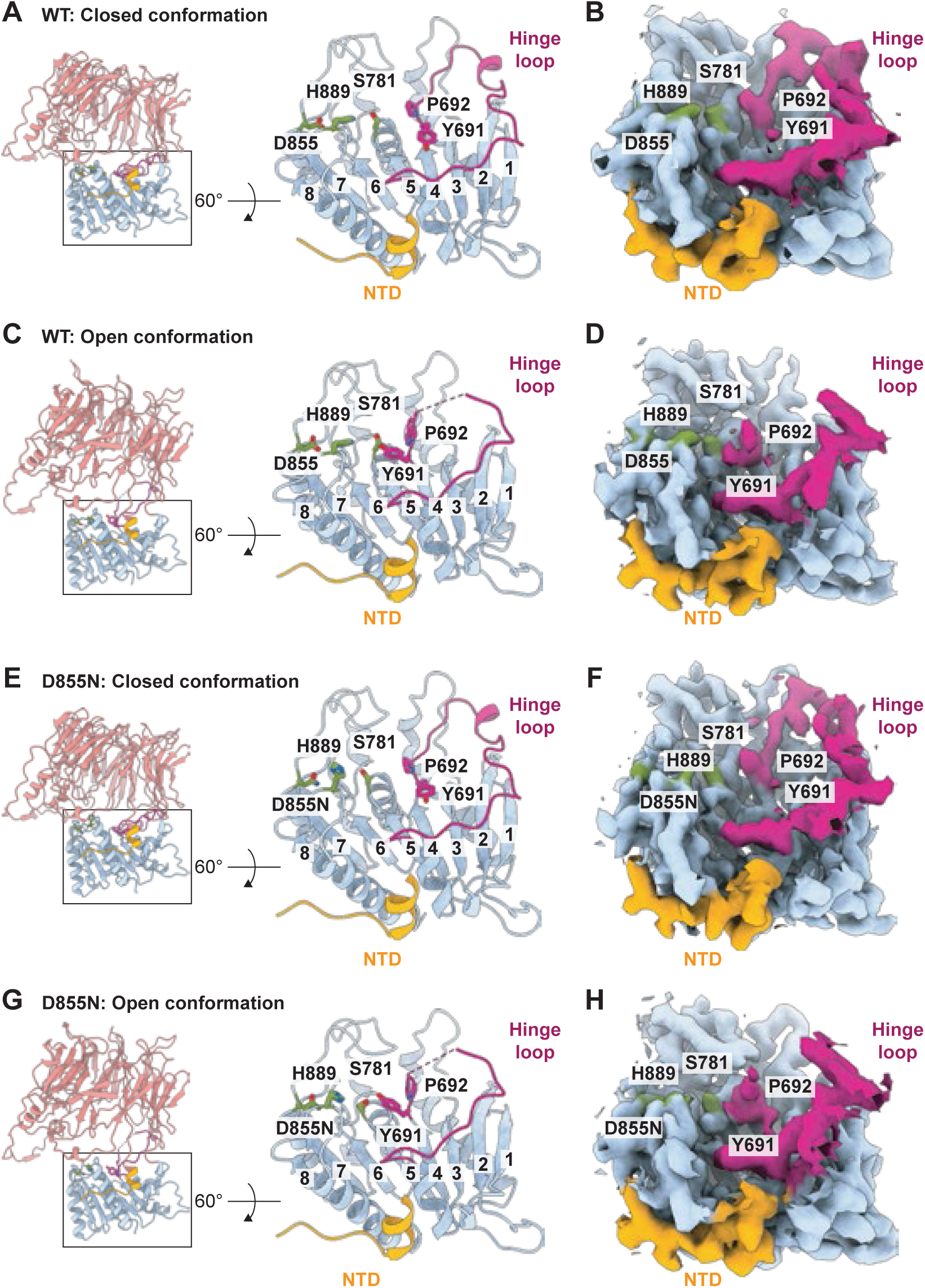
The catalytic triad and hinge loop of the wild type and D855N. (A–D) Wild type CGEP catalytic domains viewed from the top. Cartoon representation (A and C) and cryo-EM maps (B and D) are shown. Designated residues are in stick representation, and β-strands are numbered, with strand eight representing the sticky β-edge. Closed (A and B) and open (C and D) conformations are shown. (E–H) CGEP D855N catalytic domain viewed from the top. Cartoon representation (E and G) and cryo-EM maps (F and H) are shown. Designated residues are in stick representation, and β-strands are numbered, with strand eight representing the sticky β-edge. Closed (E and F) and open (G and H) conformations are shown.

To further validate this mechanism, we generated a different catalytically inactive mutant, D855N, which preserves the catalytic serine residue S781 (Fig. S8-11 and Table S4). Similar to S781R and wild⍰type CGEP, the D855N mutant also exhibited limited substrate access in the open conformation, while the catalytic triad remained essentially unchanged between the two states (Fig. 5E–H). Together, our cryo⍰EM structures strongly support a model in which CGEP is catalytically active only in the closed conformation. In the open state, the hinge loop repositions to obstruct access to the catalytic triad, thereby preventing substrate binding and subsequent catalysis.

### Hinge loop also plays a key role in substrate selectivity

CGEP preferentially cleaves peptides after a glutamate residue, largely independent of the surrounding amino acid sequence. To understand how this substrate selectivity is achieved, we examined the protein surface of the catalytic domain and identified a prominent pocket adjacent to the catalytic S781 that may serve as a selectivity cavity accommodating the glutamate side chain (Fig. 6A). Molecular docking using a portion of the CGEP CTD sequence (Asn–Pro–Glu–Phe–Gly) supports this model, showing that the glutamate side chain fits into this pocket, which is bordered by the hinge loop (Fig. 6A). This region of the hinge loop is highly conserved among plant S9D proteases (Fig. 6B), further supporting its potential key role in determining substrate specificity.

**Figure 6.**
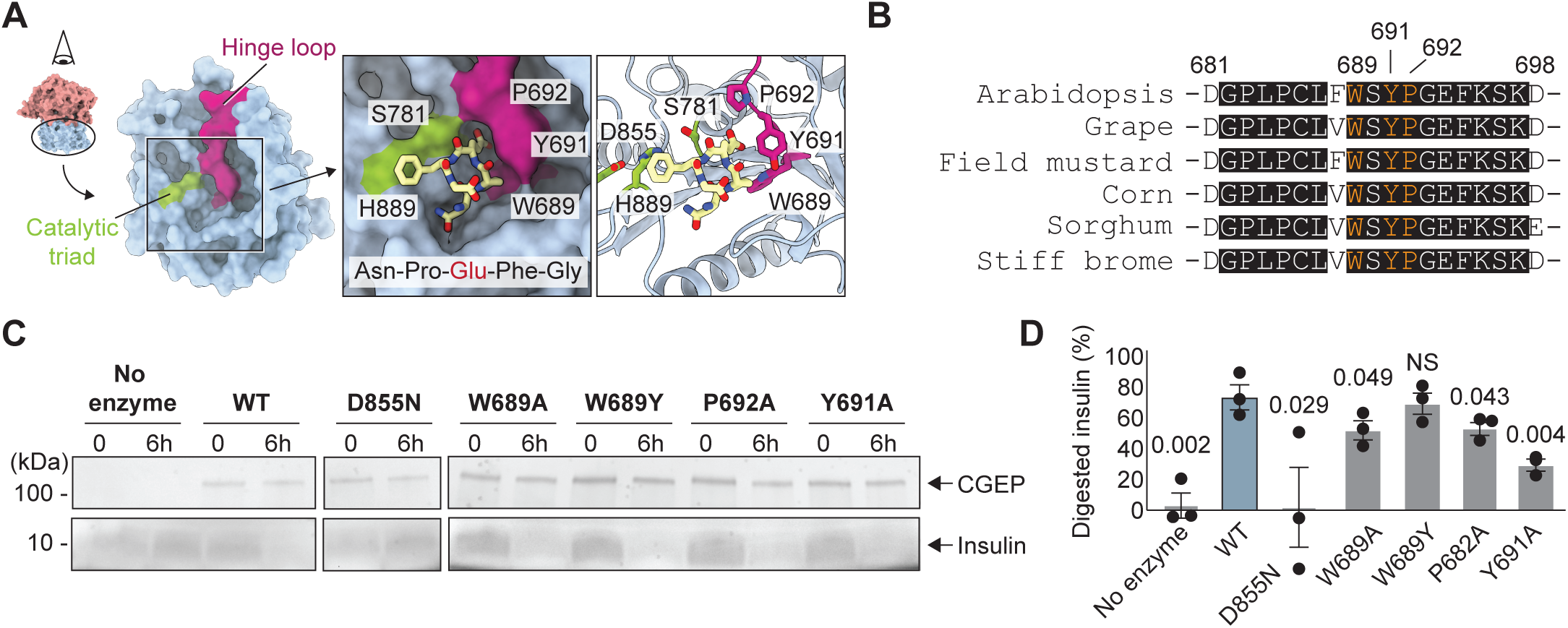
Hinge loop forms a pocket favorable for glutamate selection in the closed conformation. (A) Surface representation of the wild type CGEP catalytic domain in the closed conformation (left). Docked C-terminal peptide around the catalytic triad (Asn–Pro–Glu–Phe–Gly) using SwissDock with default settings (Bugnon et al. 2024; Grosdidier et al. 2011; Rohrig et al. 2023; Zoete et al. 2016) is shown in yellow (right). (B) Sequence alignment of hinge loops from 6 different plant (angiosperm) species (3 dicots and 3 monocots). Fully conserved residues are shaded, and residues targeted for mutagenesis are highlighted in orange. Numbers correspond to Arabidopsis residue positions. (C) SDS-PAGE of recombinant CGEP mutants (top) and insulin incubated for 6 hours (bottom). (D) Quantification of SDS-PAGE band intensity. Error bars represent standard deviation; P-values from post hoc Dunnett’s test following one-way ANOVA are shown. NS, non-significant; N = 3.

**Figure 7.**
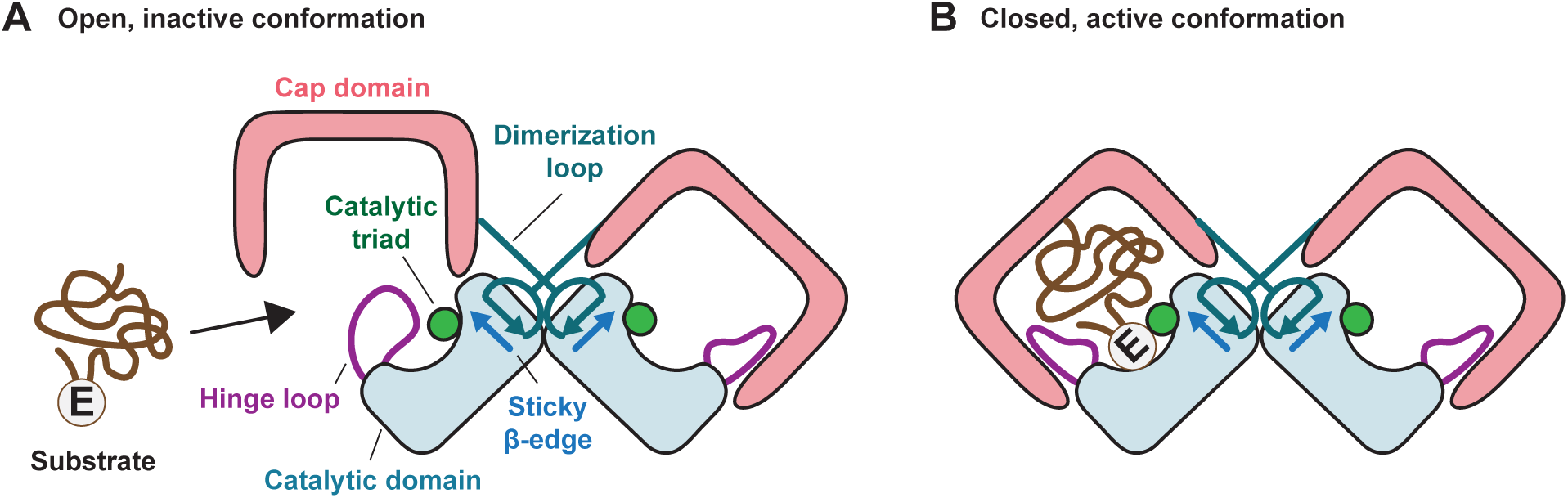
Proposed model for the S9D-specific hinge loop–mediated substrate selection mechanism. (A) A substrate containing a glutamate residue and with a size of up to 25 kDa enters the cap domain through the large side opening. In the open conformation, the hinge loop prevents the substrate from accessing the catalytic triad. (B) Upon cap domain closure, the hinge loop undergoes a conformational rearrangement, forming a pocket that favors the glutamate residue in the substrate, which is then degraded by the catalytic triad.

To investigate the functional contribution of the hinge loop, we generated W689A, W689Y, Y691A, and P692A CGEP mutant constructs, which were expressed and purified from *E. coli*. We then assessed their catalytic activity using insulin as a model substrate. Incubation of wild⍰type CGEP with insulin for six hours resulted in ∼70% substrate digestion, whereas the catalytically inactive D855N mutant showed no detectable activity (Fig. 6C, D). Under the same conditions, the conservative W689Y mutation retained activity comparable to the wild type, suggesting that this mutation does not impair substrate recognition. In contrast, more disruptive mutations—W689A, Y691A, and P692A—substantially reduced enzymatic activity. Importantly, all CGEP variants expressed well and were monodisperse and dimeric (Fig. S12), indicating that their reduced activity was not due to misfolding. Together, these results support a model in which the hinge loop forms a glutamate⍰selective pocket near the catalytic triad in the closed conformation, thereby playing a critical role in defining CGEP’s substrate preference.

## Discussion

Despite the availability of high-resolution structures for S9A, B and C proteases, mechanistic insights into the S9D family have remained limited due to the absence of resolved structures and due to its poor sequence similarity with other subtypes. In this study, we determined the first high-resolution structures of an S9D family member, CGEP, expressed in both plants and bacteria. As predicted, CGEP consists of two major domains: a β-propeller cap and an α/β/α sandwich catalytic domain. We found that CGEP predominantly exists as a dimer, similar to S9B proteases, but exhibits multiple structural features unique to the S9D family. These features likely underlie S9D-specific characteristics, particularly its strong preference for cleaving after glutamate residues. Based on our findings, we propose that the hinge loop functions as a gate that restricts catalytic activity to the closed conformation. Our proposed mechanism is summarized in Figure 8.

Unlike other dimeric S9 proteases, CGEP dimerization is stabilized by two distinct mechanisms. The first interface is mediated by hydrophobic interactions between the catalytic domains, while the second involves interdomain β-sheet formation between the cap and catalytic domains. These additional interdomain interactions likely reinforce the dimeric core, providing a rigid scaffold that supports the conformational rearrangement of the hinge loop during cap opening and closing. Furthermore, the interdomain β-sheet may shield the sticky β-edge, which has been implicated in aggregation (Kiss-Szeman et al. 2019).

Another unique feature of CGEP is that its open conformation is spacious enough to accommodate substrates up to ∼20 kDa (or ∼40 Å in diameter), yet substrate access to the catalytic triad is sterically restricted by the hinge loop. CGEP preserves the integrity of its catalytic triad in both conformations, relying instead on hinge-loop gating to control activity. This regulatory mechanism differs from other S9 proteases that also undergo open–close transitions, where cap opening disrupts the catalytic triad—such as displacement of the catalytic histidine—rendering the enzyme inactive. For S9 proteases with a static catalytic triad, substrate size selectivity is achieved through an inherently narrow access pathway to the active site (Menyhard et al. 2013; Rasmussen et al. 2003). This contrasts with CGEP, which uses an open conformation to accommodate larger substrates, such as insulin (10 kDa), while maintaining catalytic regulation through hinge-loop positioning.

Our molecular docking and mutagenesis studies suggest a potential mechanism underlying the glutamate-specific protease activity of S9D family enzymes. During cap closure, the hinge loop undergoes a substantial conformational rearrangement, forming a pocket that favors glutamate side chains near the catalytic triad. Notably, alanine substitutions of the bulky side chains at W689 and Y691 significantly reduce CGEP’s ability to digest insulin, rather than increasing activity as might be expected if hinge-loop flexibility were the limiting factor for substrate access. These findings support the idea that, in its closed conformation, the hinge loop, together with neighboring residues of the catalytic triad, creates a microenvironment that facilitates glutamate binding. Nevertheless, these results alone cannot provide a definitive conclusion. Although we attempted to visualize the substrate-binding site using a catalytically inactive D855N mutant supplemented with synthetic glutamate-containing peptides, the cryo-EM density of the trapped substrate was too weak to draw definitive conclusions. Determining high-resolution structures of mutants such as W689A or Y691A, which exhibit diminished catalytic activity likely due to compromised glutamate binding, will be helpful to further elucidate the structural basis of substrate selectivity.

We attempted to visualize the CTD in our cryo-EM experiments by mutating the catalytic serine residue to arginine (S781R) in CGEP transgenically expressed in *Arabidopsis*. However, the cryo-EM density for the C-terminal region downstream of N915 (total of 71 AA), including the autocleavage site E946, was poorly resolved. This is perhaps not surprising since the predicted structure by AlphaFold shows a random coil after residue 912.

In conclusion, we present the first high-resolution structures of a plant S9D protease, CGEP, uncovering a regulatory mechanism distinct from other S9 proteases. Most notably, the discovery that hinge-loop dynamics govern substrate access and selectivity sets the S9D subfamily apart. These findings establish a new paradigm for S9D protease regulation and substrate recognition, providing a structural foundation for future mechanistic and functional studies.

## Materials and methods

### CGEP purification from transgenic *Arabidopsis*

A stable transgenic line of catalytically inactive CGEP S781R with a C-terminal StrepII tag (Bhuiyan et al. 2020) was used to purify the CGEP mutant from transgenic *Arabidopsis*. The StrepII protein purification was performed as described (Rei Liao et al. 2022). The purified protein was buffer exchanged into 50 mM HEPES-KOH pH 8.0, 75 mM NaCl, and 10 mM MgCl_2_ concentrated on 30 kDa centrifugal filters. Protein sample purity was assessed by SDS-PAGE.

### Molecular cloning and recombinant protein purification

His tags were added to the N-terminus of the AtCGEP coding sequence (starting from because the cTP was removed) with GG linkers, and they were subcloned into pET28 vectors using standard molecular cloning procedures. The SMILES string of the peptide with the c-terminal sequence “NPEFG” was docked into the closed conformation of a CGEP monomer using SwissDock (Bugnon, 2024) with default parameters surrounding the catalytic triad. Candidate residues in proximity to the peptide binding pose were chosen for site directed mutagenesis using quickchange PCR.

BL21 cells harboring CGEP constructs were grown at 37°C with 250 rpm shaking in 50 mL Luria Broth overnight. 500 mL Terrific Broth cultures were inoculated with the starter culture and allowed to grow for ∼3 hours at 37°C with 350 rpm shaking. Expression was induced in cultures at an OD between 2.0 – 4.0 using 1 mM IPTG. The temperature was lowered to 22°C and cells expressed CGEP proteins for 3 hours before harvest by centrifugation. Cells were washed once with PBS and flash frozen.

Frozen cell pellets were thawed in 30 mL lysis buffer (50 mM Tris HCl pH 8.0, 500 mM NaCl, 10% glycerol) plus 0.5 mg/ml lysozyme, 0.1 mg/ml DNase, and 10 mM MgCl_2_ at 4°C with shaking. Cells were further lysed with multiple rounds of sonication until the lysate was less viscous and darker in color.

The lysate was cleared by ultracentrifugation, then proteins were bound to equilibrated TALON superflow resin (Cytiva Life Sciences; cat. 2895702) by tipping at 4°C for 2 hours. Resin was washed with 50 column volumes of lysis buffer plus 5 mM imidazole pH 8.0. Protein was eluted by lysis buffer containing 400 mM imidazole pH 8.0.

Eluent was concentrated to 0.5 mL using an Amicon spin column with a 50k cutoff (Millipore; cat. UFC9050). Protein was subjected to size exclusion chromatography on a superdex 200 increase 10/300 GL column (Cytiva Life Sciences; cat. 28990944) with 100 mM NaCl and 50 mM Tris HCl (pH 8) running buffer. Fractions were pooled and prepared for either cryo-EM or substrate digestion assays.

### Substrate digestion assay

Substrate digestion assays were carried out as described before with minor modifications (Bhuiyan et al. 2020). Briefly, glycerol was added to pooled SEC fractions to a final concentration of 10%. The identity and purity of the collected factions were assessed by SDS-PAGE followed by Oriole staining (Bio-Rad). Protein concentrations were determined by Nanodrop. Purified CGEP was mixed at a final concentration of 50 µg/ml and 25 µg/ml insulin (Alfa Aesar, #J67626) in 0.1 mL reactions in 100 mM NaCl, 50 mM Tris-HCL (pH 8) and 10% glycerol and incubated for 6 hours at 37°C. Reactions were stopped by adding SDS to a final concentration of 3%. Efficiency of substrate digestion was assessed on a 4-20% gradient reducing SDS-PAGE gel (Biorad, #4561094) and stained with Oriole fluorescent gel (Biorad, #161-0496) stain for 1 hour. Band intensities from three individual replicates were estimated using ImageJ and averaged. A one-way ANOVA test followed by a two-tailed T-test was used to compare each mutant to the WT.

### Cryo-EM sample preparation and data collection

Protein samples were spotted on Quantifoil 1.2/1.3 holey carbon copper grids with 400 mesh (Electron Microscopy Services; cat. Q450CR1.3). 3 µL of protein were blotted for 4 seconds with force 7 on an FEI Mark IV Vitrobot with humidity set to 100% and chamber temperature set to 4°C. Samples were plunge frozen into liquid ethane and stored in liquid nitrogen.

Images of CGEP protein produced in Arabidopsis were collected on the Talos Arctica with K3 direct electron detector with a GIF Bioquantum energy filter from Cornell Center for Materials Resources. Nominal magnification was 63,000x, resulting in the superresoution pixel size of 0.685 Å/pix. The total electron dose was 40 e^-^/Å/pix over 40 frames. Images of recombinantly produced CGEP were taken on the Titan Krios with a Falcon 4i direct electron detector and GIF Biocontinuum energy filter at Pacific Northwest Cryo-EM Center. Nominal magnification was 96,000x, and after upsampling with a factor of 2, the pixel size was 0.82 Å/pix. The total electron dose was 50 e^-^/Å/pix.

### Cryo-EM data processing and atomic model building

Cryo-EM data processing for each protein is summarized in supplemental figures S3, S8, and S9. Briefly, the movies were motion-corrected and contrast transfer function (CTF) estimated. Micrographs of sufficient ice thickness were subject to blob picker for an 80-120 Å blob. Picks were extracted and 2D classified, then the well-ordered templates were used for template picker. Template picked particles were extracted and 2D classified to remove non-protein particles. A small number of particles were subject to *ab initio* map generation. The most promising maps were used in heterogenous refinement with all classified particles. Iterative rounds of heterogenous refinement, 3D classification, and non-uniform refinement sorted out both low-resolution particles and multiple conformations in the dataset. Final particle stacks were assembled into 3D maps using non-uniform refinement with CTF parameters enabled. Maps were manually sharpened according to the b-factor that enhanced the features but did not amplify noise. For applicable maps, C2 symmetry was applied at this step only. All processing jobs were carried out in cryoSPARC v4.7.0. The quality of the cryo-EM maps is summarized in supplemental figures S4 and S10.

The Alphafold prediction for CGEP was fit into one subunit of the closed-closed map, then manually adjusted to fit the density. NCS parameters were generated for the map in Phenix, then applied to the atomic model to generate a dimeric structure. For the open-open maps, one subunit of the closed conformation was fit into the density and adjusted accordingly. The same NCS protocol was employed to generate dimeric structures. For the closed-open maps, one protomer from the closed-closed model and one protomer from the open-open model were combined and fit to the density. All models were refined using iterative real-space refinement in Phenix until the parameters stopped improving. Map to model fitness is summarized by Q-scores and model vs map FSCs in figures S5 and S11.

## Acknowledgements

We thank members of the Kawate and Van Wijk labs helpful discussions. This work was supported by National Science Foundation grant MCB #2222495 to van Wijk and Kawate. The Kawate lab was supported by National Institutes of Health grant R01GM114379 and Cornell CVM Graduate Scholarship. This work made use of the Cornell Center for Materials Research shared instrumentation facility. This work relied on data collected using an instrument supported by the NIH through award S10OD030470. A portion of this research was supported by NIH grant R24GM154185 and performed at the Pacific Northwest Center for Cryo-EM (PNCC) with assistance from James Evans, Kjirsten Wheeler, Rose Marie Yasuda, Sean Mulligan, Marzia Miletto, and Nancy Meyer.

## Data sharing plan

Cryo-EM maps are deposited to the Electron Microscopy Data Bank (EMDB) and atomic models are deposited to the Protein Data Bank (PDB) under the following ascension numbers: structures purified from *Arabidopsis*: closed-closed (EMD-75113/ 10EO), closed-open (EMD-75114/ 10EP), open-open (EMD-75116/ 10ER). Structures purified from *E. coli*: WT structures – closed-open (EMD-75115/ 10EQ), open-open (EMD-75117/ 10ES). Recombinant D855N structures – closed-open (EMD-75118/ 10ET), open-open (EMD-75119/ 10EU). Motion corrected datasets are deposited to the Electron Microscopy Public Image Archive under the following ascension numbers: Arabidopsis dataset (EMPIAR-13207). Recombinant WT dataset (EMPIAR-13208). Recombinant D855N dataset (EMPIAR-13206). All other data to support these findings is found within this manuscript or is available upon reasonable request to the authors.

## Abbreviations and symbols

CGEP: chloroplast glutamyl endopeptidase
Cryo-EM: cryo-electron microscopy
POP: Prolyl oligopeptidase
DPP-IV: Dipeptidyl peptidase IV
AAP: Acylaminoacyl peptidase

## Notes

### Competing Interest Statement

The authors have declared no competing interest.

