## Supplementary materials for "Structural basis for dimerization, catalytic regulation, and substrate selectivity in S9D proteases"

Figure S1

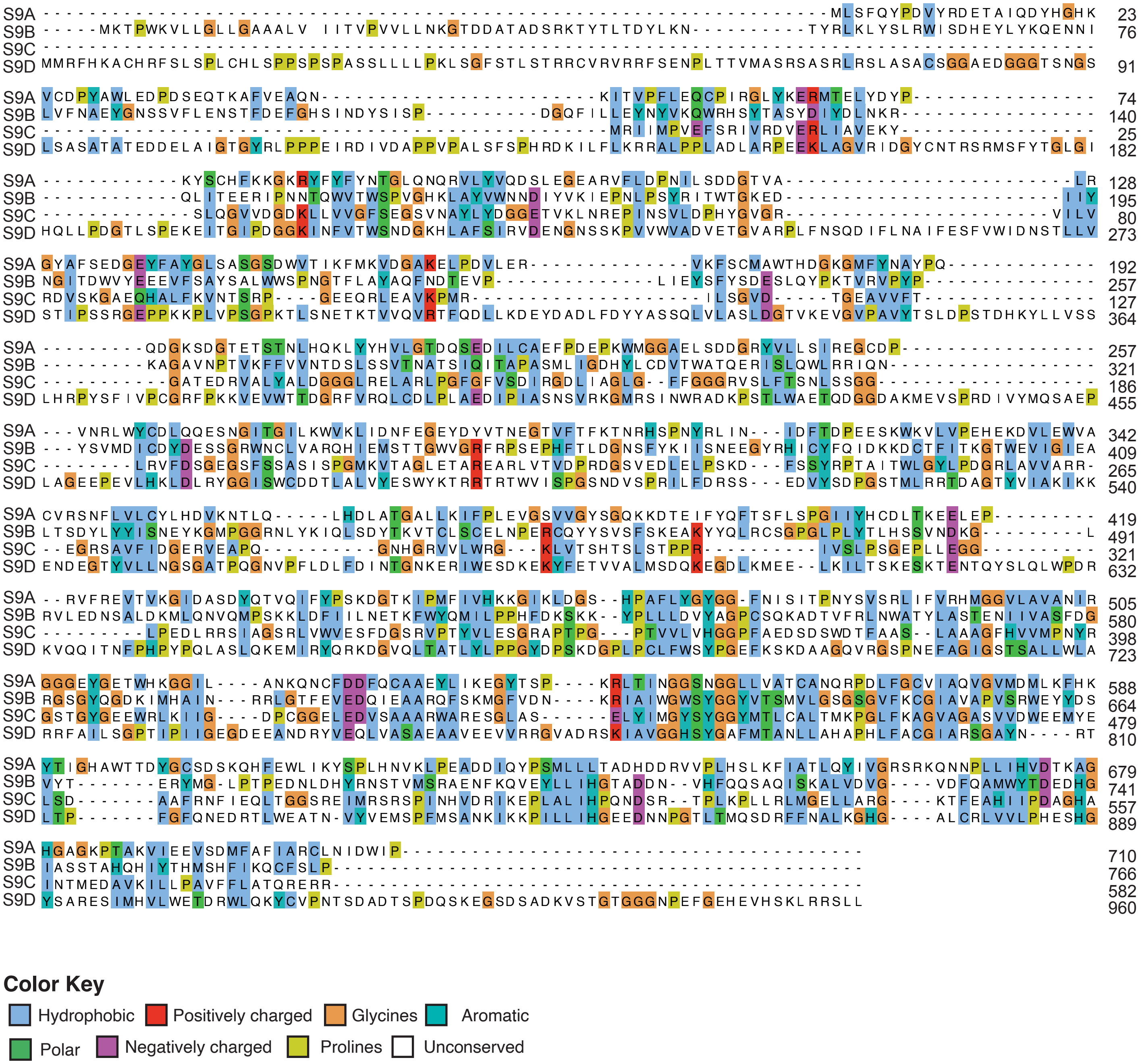
S9A --------------------------------------------------------------------MLSFQYPDVYRDETAIQDYHGHK 23
S9B -----MKTPWKVLLGLLGAAALV IITVPVVLLNKGTDDATADSRKTYTLTDYLKN----------TYRLKLYSLRWISDHEYLYKQENNI 76
S9C -------------------------------------------------------------------------------------------
S9D MMRFHKACHRFSLSPLCHLSPPSPSPASSLLLLPKLSGFSTLSTRRCVRVRRFSENPLTTVMASRSASRLRSLASACSGGAEDGGGTSNGS 91
S9A VCDPYAWLEDPDSEQTKAFVEAQN------------------------KITVPFLEQCPIRGLYKERMTELYDYP---------------- 74
S9B LVFNAEYGNSSVFLENSTFDEFGHSINDYSISP----------DGQFILLEYNYVKQWRHSYTASYDIYDLNKR----------------- 140
S9C -------------------------------------------------MRIIMPVEFSRIVRDVERLIAVEKY----------------- 25
S9D LSASATATEDDELAIGTGYRLPPPEIRDIVDAPPVPALSFSPHRDKILFLKRRALPPLADLARPEEKLAGVRIDGYCNTRSRMSFYTGLGI 182
S9A ------------KYSCHFKKGKRYFYFYNTGLQNQRVLYVQDSLEGEARVFLDPNILSDDGTVA-------------------------LR 128
S9B ------------QLITEERIPNNTQWVTWSPVGHKLAYVWNNDIYVKIEPNLPSYRITWTGKED------------------------IIY 195
S9C -------------SLQGVVDGDKLLVVGFSEGSVNAYLYDGGETVKLNREPINSVLDPHYGVGR-----------------------VILV 80
S9D HQLLPDGTLSPEKEITGIPDGGKINFVTWSNDGKHLAFSIRVDENGNSSKPVVWVADVETGVARPLFNSQDIFLNAIFESFVWIDNSTLLV 273
S9A GYAFSEDGEYFAYGLSASGSDWVTIKFMKVDGAKELPDVLER--------------VKFSCMAWTHDGKGMFYNAYPQ------------- 192
S9B NGITDWVYEEEVFSAYSALWWSPNGTFLAYAQFNDTEVP--------------LIEYSFYSDESLQYPKTVRVPYP--------------- 257
S9C RDVSKGAEQHALFKVNTSRP----GEEQRLEAVKPMR--------------------ILSGVD-----TGEAVVFT--------------- 127
S9D STIPSSRGEPPKKPLVPSGPKTLSNETKTVVQVRTFQDLLKDEYDADLFDYYASSQLVLASLDGTVKEVGVPAVYTSLDPSTDHKYLLVSS 364
S9A ---------QDGKSDGTETSTNLHQKLYYHVLGTDQSEDILCAEFPDEPKWMGGAELSDDGRYVLLSIREGCDP----------------- 257
S9B ---------KAGAVNPTVKFFVVNTDSLSSVTNATSIQITAPASMLIGDHYLCDVTWATQERISLQWLRRIQN------------------ 321
S9C -----------GATEDRVALYALDGGGLRELARLPGFGFVSDIRGDLIAGLG--FFGGGRVSLFTSNLSSGG------------------- 186
S9D LHRPYSFIVPCGRFPKKVEVWTTDGRFVRQLCDLPLAEDIPIASNSVRKGMRSINWRADKPSTLWAETQDGGDAKMEVSPRDIVYMQSAEP 455
S9A ---VNRLWYCDLQQESNGITGILKWVKLIDNFEGEYDYVTNEGTVFTFKTNRHSPNYRLIN---IDFTDPEESKWKVLVPEHEKDVLEWVA 342
S9B ---YSVMDICDYDESSGRWNCLVARQHIEMSTTGWVGRFRPSEPHFTLDGNSFYKIISNEEGYRHICYFQIDKKDCTFITKGTWEVIGIEA 409
S9C --------LRVFDSGEGSFSSASISPGMKVTAGLETAREARLVTVDPRDGSVEDLELPSKD---FSSYRPTAITWLGYLPDGRLAVVARR- 265
S9D LAGEEPEVLHKLDLRYGGISWCDDTLALVYESWYKTRRTRTWVISPGSNDVSPRILFDRSS---EDVYSDPGSTMLRRTDAGTYVIAKIKK 540
S9A CVRSNFLVLCYLHDVKNTLQ-----LHDLATGALLKIFPLEVGSVVGYSGQKKDTEIFYQFTSFLSPGIIYHCDLTKEELEP--------- 419
S9B LTSDYLYYISNEYKGMPGGRNLYKIQLSDYTKVTCLSCELNPERCQYYSVSFSKEAKYYQLRCSGPGLPLYTLHSSVNDKG---------L 491
S9C --EGRSAVFIDGERVEAPQ----------GNHGRVVLWRG---KLVTSHTSLSTPPR----------IVSLPSGEPLLEGG---------- 321
S9D ENDEGTYVLLNGSGATPQGNVPFLDLFDINTGNKERIWESDKEKYFETVVALMSDQKEGDLKMEE--LKILTSKESKTENTQYSLQLWPDR 632
S9A --RVFREVTVKGIDASDYQTVQIFYPSKDGTKIPMFIVHKKGIKLDGS--HPAFLYGYGG-FNISITPNYSVSRLIFVRHMGGVLAVANIR 505
S9B RVLEDNSALDKMLQNVQMPSKKLDFIILNETKFWYQMILPPHFDKSKK--YPLLLDVYAGPCSQKADTVFRLNWATYLASTENIIVASFDG 580
S9C ---------LPEDLRRSIAGSRLVWVESFDGSRVPTYVLESGRAPTPG---PTVVLVHGGPFAEDSDSWDTFAAS--LAAAGFHVVMPNYR 398
S9D KVQQITNFPHPYPQLASLQKEMIRYQRKDGVQLTATLYLPPGYDPSKDGPLPCLFWSYPGEFKSKDAAGQVRGSPNEFAGIGSTSALLWLA 723
S9A GGGEYGETWHKGGIL----ANKQNCFDDFQCAAEYLIKEGYTSP----KRLTINGGSNGGLLVATCANQRPDLFGCVIAQVGVMDMLKFHK 588
S9B RGSGYQGDKIMHAIN---RRLGTFEVEDQIEAARQFSKMGFVDN----KRIAIWGWSYGGYVTSMVLGSGSGVFKCGIAVAPVSRWEYYDS 664
S9C GSTGYGEEWRLKIIG----DPCGGELEDVSAAARWARESGLAS------ELYIMGYSYGGYMTLCALTMKPGLFKAGVAGASVVDWEEMYE 479
S9D RRFAILSGPTIPIIGEGDEEANDRYVEQLVASAEAAVEEVVRRGVADRSKIAVGGHSYGAFMTANLLAHAPHLFACGIARSGAYN----RT 810
S9A YTIGHAWTTDYGCSDSKQHFEWLIKYSPLHNVKLPEADDIQYPSMLLLTADHDDRVVPLHSLKFIATLQYIVGRSRKQNNPLLIHVDTKAG 679
S9B VYT-------ERYMG-LPTPEDNLDHYRNSTVMSRAENFKQVEYLLIHGTADDN--VHFQQSAQISKALVDVG----VDFQAMWYTDEDHG 741
S9C LSD-------AAFRNFIEQLTGGSREIMRSRSPINHVDRIKEPLALIHPQNDSR--TPLKPLLRLMGELLARG----KTFEAHIIPDAGHA 557
S9D LTP-------FGFQNEDRTLWEATN-VYVEMSPFMSANKIKKPILLIHGEEDNNPGTLTMQSDRFFNALKGHG----ALCRLVVLPHESHG 889
S9A HGAGKPTAKVIEEVSDMFAFIARCLNIDWIP---------------------------------------- 710
S9B IASSTAHQHIYTHMSHFIKQCFSLP---------------------------------------------- 766
S9C INTMEDAVKILLPAVFFLATQRERR---------------------------------------------- 582
S9D YSARESIMHVLWETDRWLQKYCVPNTSDADTSPDQSKEGSDSADKVSTGTGGGNPEFGEHEVHSKLRRSLL 960
Color Key
Hydrophobic
Positively charged
Glycines
Aromatic
Polar
Negatively charged
Prolines
Unconserved

Figure S2

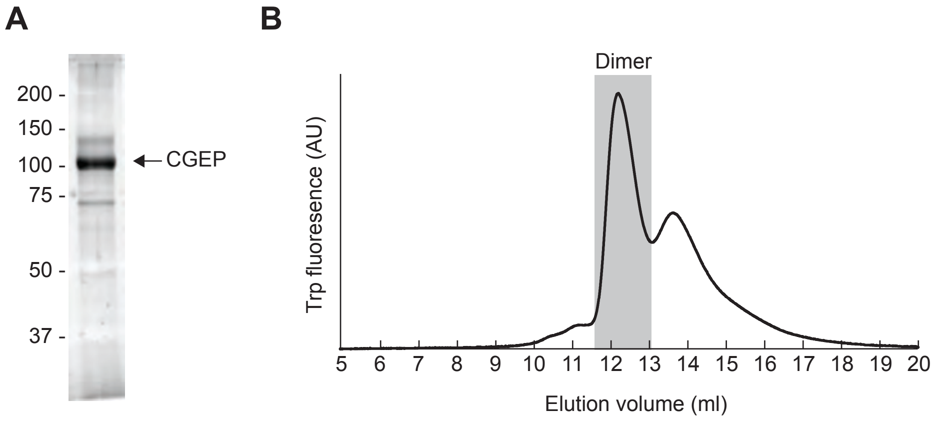
A
200 -
150 -
100 -
75 -
50 -
37 -
CGEP
B
Dimer
Trp fluoresence (AU)
5 6 7 8 9 10 11 12 13 14 15 16 17 18 19 20
Elution volume (ml)

Figure S3

**CGEP S781R**

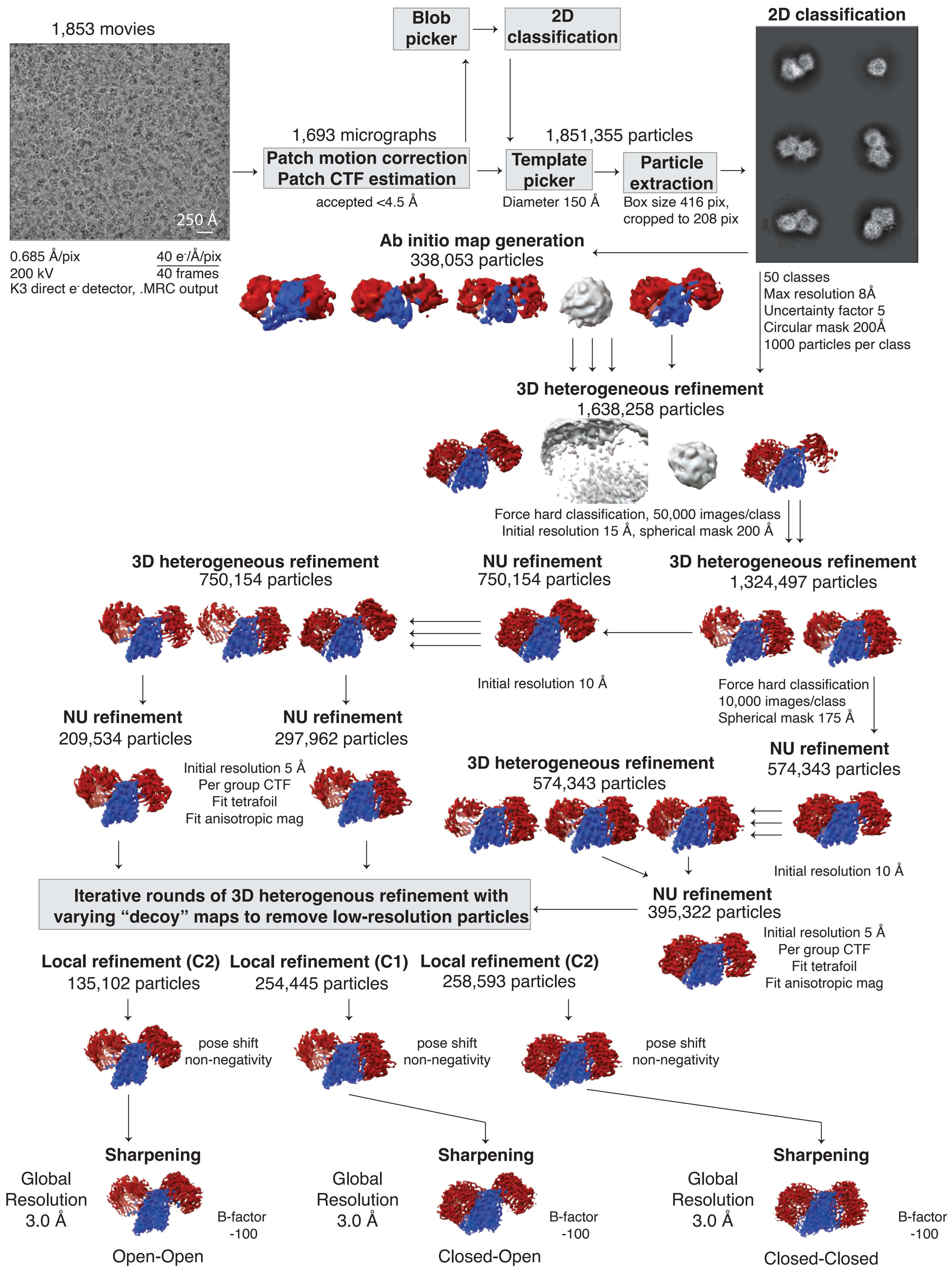

Figure S4

**Cryo-EM map validation of CGEP S781R** EMPIAR-13207

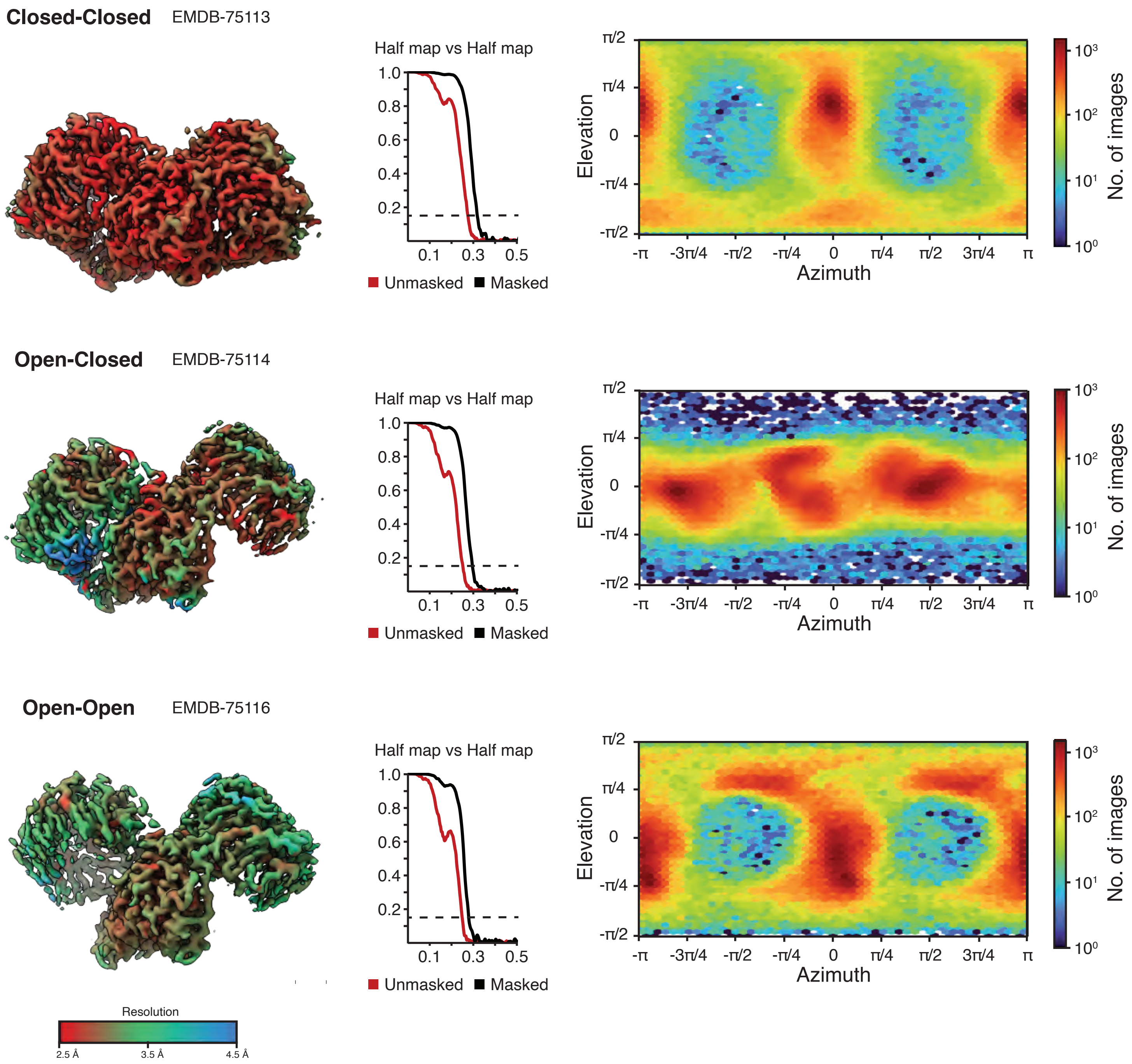

Figure S5

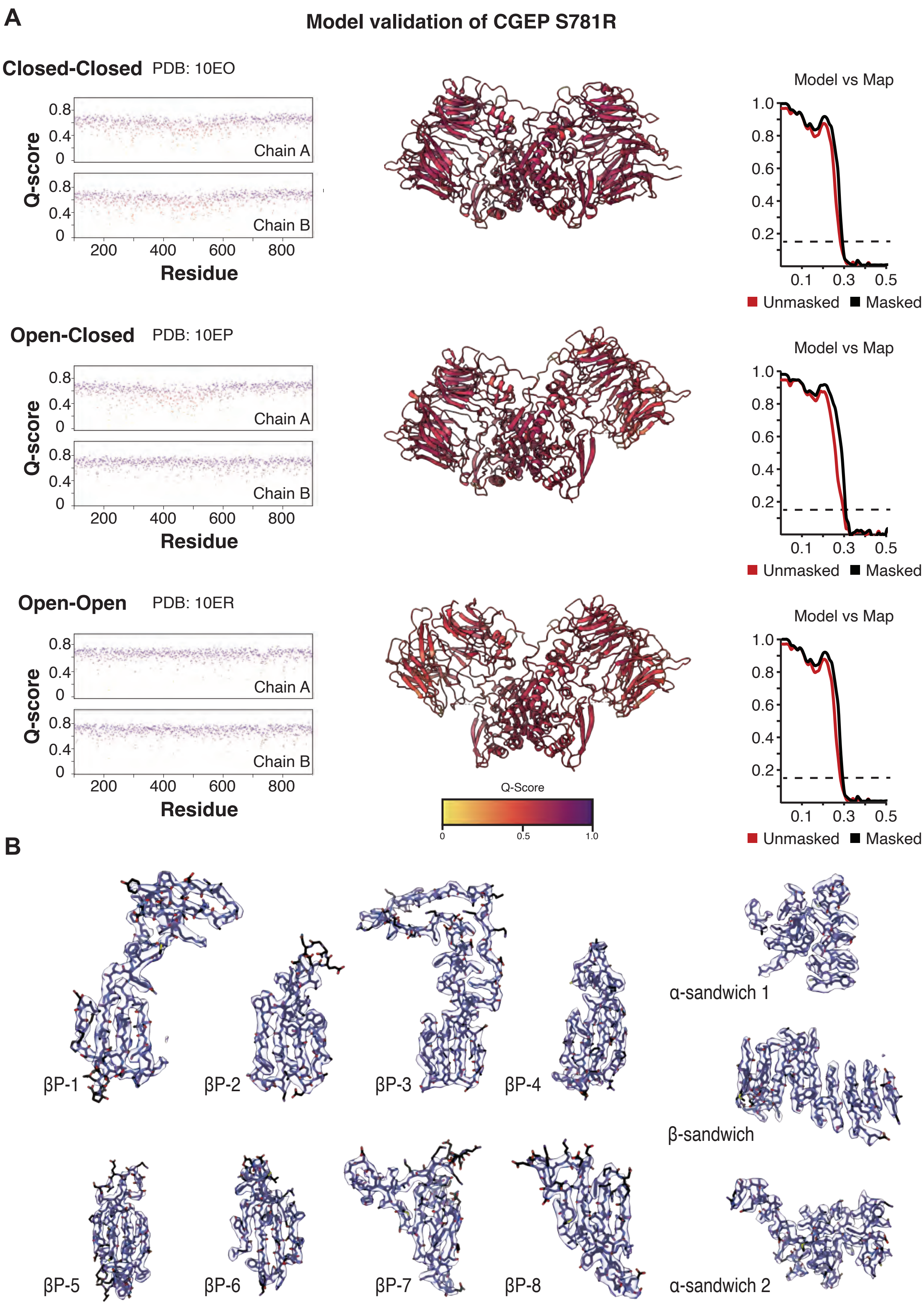
A
Model validation of CGEP S781R
Closed-Closed PDB: 10EO
Q-score
0.8
0.4
0
Chain A
Chain B
200
400
600
800
Residue
Model vs Map
1.0
0.8
0.6
0.4
0.2
0.1
0.3
0.5
Unmasked
Masked
Open-Closed PDB: 10EP
Open-Open PDB: 10ER
Q-Score
0
0.5
1.0
B
βP-1
βP-2
βP-3
βP-4
α-sandwich 1
β-sandwich
βP-5
βP-6
βP-7
βP-8
α-sandwich 2

Figure S6

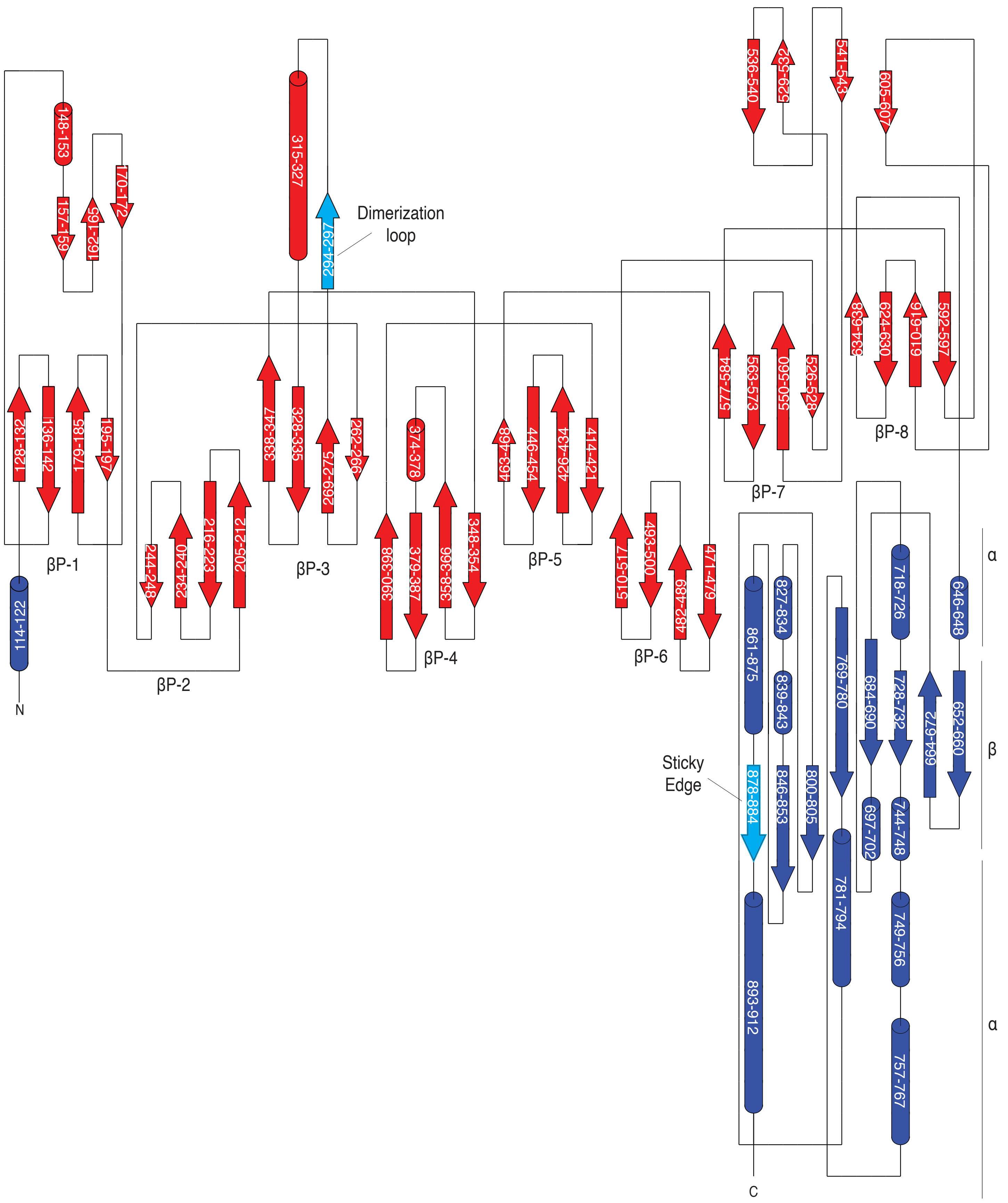

148-153
157-159
162-165
170-172
128-132
136-142
179-185
195-197
βP-1
114-122
N
244-248
234-240
216-223
205-212
βP-2
315-327
294-297
Dimerization loop
338-347
328-335
269-275
262-266
βP-3
374-378
390-398
379-387
358-366
348-354
βP-4
463-468
446-454
426-434
414-421
βP-5
510-517
493-500
482-489
471-479
βP-6
536-540
529-532
541-543
605-607
577-584
563-573
550-560
526-528
βP-7
634-638
624-630
610-616
592-597
βP-8
861-875
827-834
839-843
878-884
846-853
800-805
Sticky Edge
769-780
781-794
684-690
697-702
718-726
728-732
744-748
749-756
757-767
664-672
646-648
652-660
α
β
α
893-912
C

Figure S7

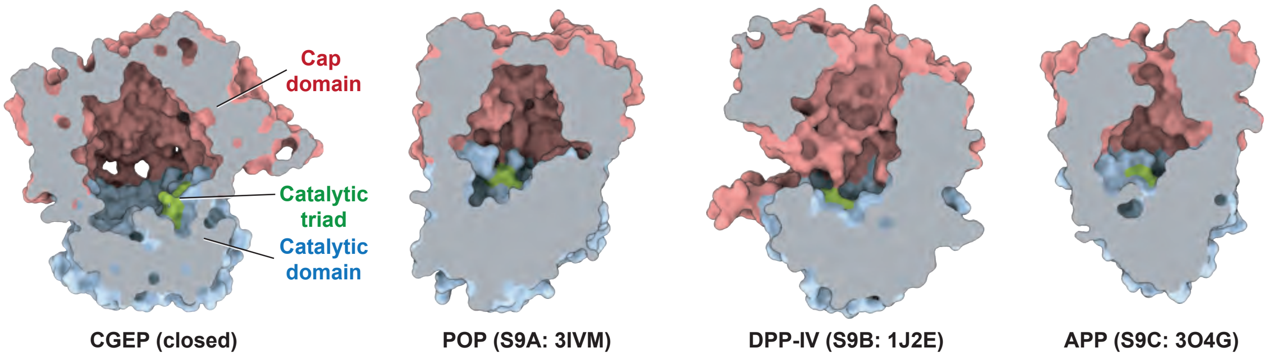
Cap
domain
Catalytic
triad
Catalytic
domain
CGEP (closed)
POP (S9A: 3IVM)
DPP-IV (S9B: 1J2E)
APP (S9C: 3O4G)

Figure S8

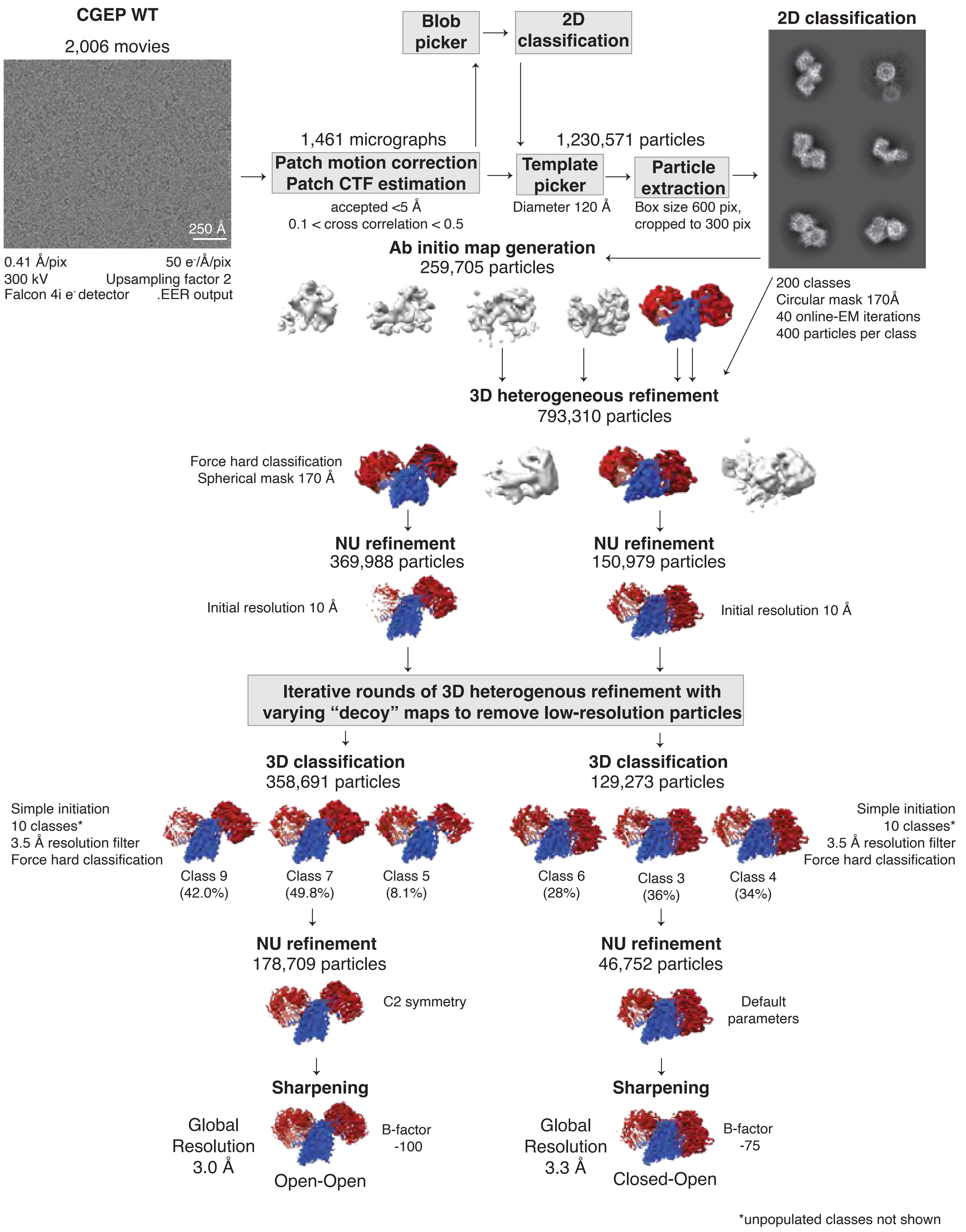

CGEP WT
2,006 movies
250 Å
0.41 Å/pix
50 e-/Å/pix
300 kV
Upsampling factor 2
Falcon 4i e- detector
.EER output
1,461 micrographs
Patch motion correction
Patch CTF estimation
accepted <5 Å
0.1 < cross correlation < 0.5
Blob picker
2D classification
1,230,571 particles
Template picker
Diameter 120 Å
Particle extraction
Box size 600 pix, cropped to 300 pix
2D classification
200 classes
Circular mask 170Å
40 online-EM iterations
400 particles per class
Ab initio map generation
259,705 particles
3D heterogeneous refinement
793,310 particles
Force hard classification
Spherical mask 170 Å
NU refinement
369,988 particles
NU refinement
150,979 particles
Initial resolution 10 Å
Initial resolution 10 Å
Iterative rounds of 3D heterogenous refinement with varying "decoy" maps to remove low-resolution particles
3D classification
358,691 particles
3D classification
129,273 particles
Simple initiation
10 classes*
3.5 Å resolution filter
Force hard classification
Class 9 (42.0%)
Class 7 (49.8%)
Class 5 (8.1%)
Class 6 (28%)
Class 3 (36%)
Class 4 (34%)
Simple initiation
10 classes*
3.5 Å resolution filter
Force hard classification
NU refinement
178,709 particles
C2 symmetry
NU refinement
46,752 particles
Default parameters
Sharpening
Global Resolution 3.0 Å
B-factor -100
Open-Open
Sharpening
Global Resolution 3.3 Å
B-factor -75
Closed-Open
*unpopulated classes not shown

Figure S9

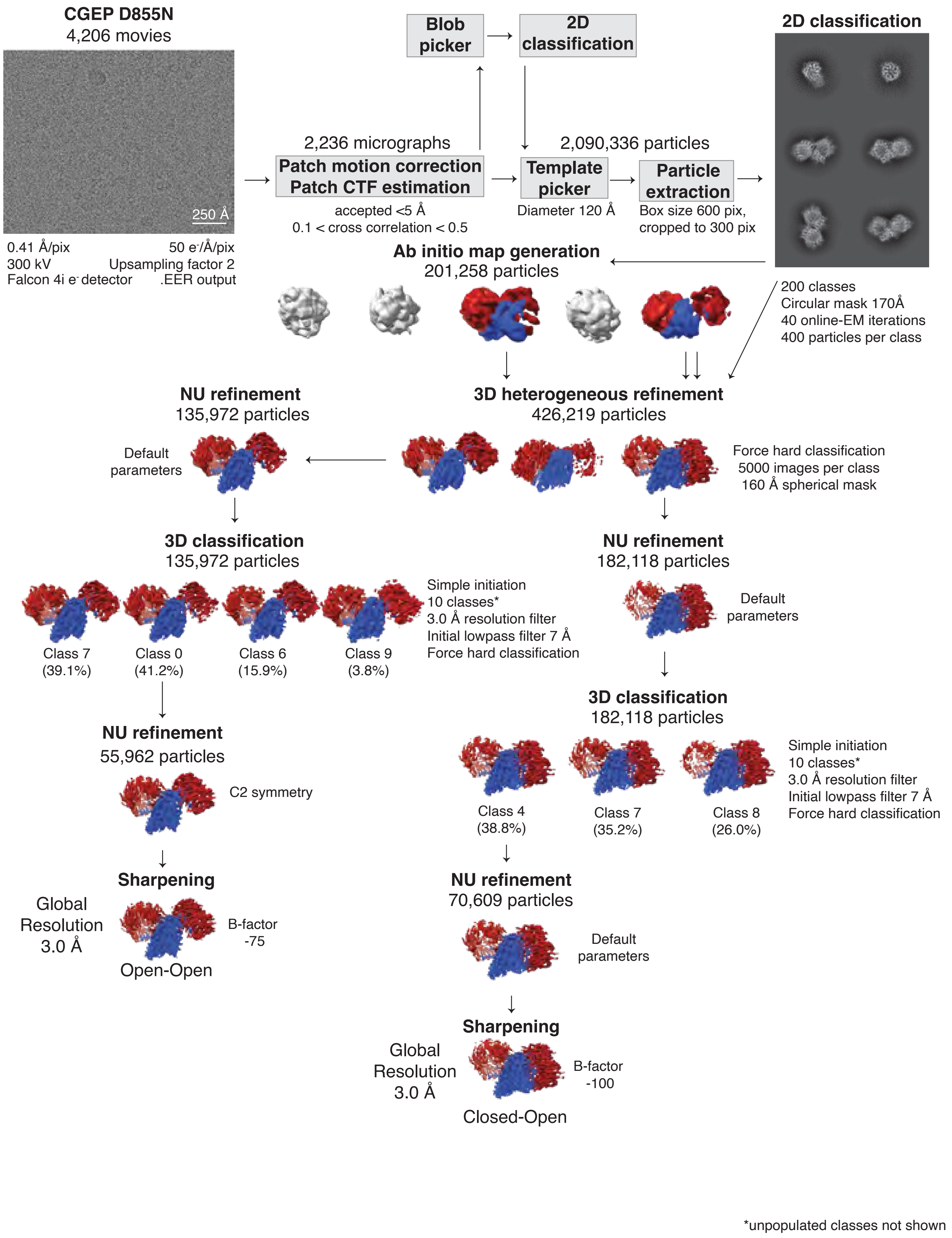

CGEP D855N
4,206 movies
250 Å
0.41 Å/pix
50 e-/Å/pix
300 kV
Upsampling factor 2
Falcon 4i e- detector
.EER output
2,236 micrographs
Patch motion correction
Patch CTF estimation
accepted <5 Å
0.1 < cross correlation < 0.5
Blob picker
2D classification
2,090,336 particles
Template picker
Diameter 120 Å
Particle extraction
Box size 600 pix, cropped to 300 pix
2D classification
200 classes
Circular mask 170Å
40 online-EM iterations
400 particles per class
Ab initio map generation
201,258 particles
3D heterogeneous refinement
426,219 particles
Force hard classification
5000 images per class
160 Å spherical mask
NU refinement
135,972 particles
Default parameters
3D classification
135,972 particles
Class 7 (39.1%)
Class 0 (41.2%)
Class 6 (15.9%)
Class 9 (3.8%)
Simple initiation
10 classes*
3.0 Å resolution filter
Initial lowpass filter 7 Å
Force hard classification
NU refinement
55,962 particles
C2 symmetry
Sharpening
Global Resolution 3.0 Å
B-factor -75
Open-Open
NU refinement
182,118 particles
Default parameters
3D classification
182,118 particles
Class 4 (38.8%)
Class 7 (35.2%)
Class 8 (26.0%)
Simple initiation
10 classes*
3.0 Å resolution filter
Initial lowpass filter 7 Å
Force hard classification
NU refinement
70,609 particles
Default parameters
Sharpening
Global Resolution 3.0 Å
B-factor -100
Closed-Open
*unpopulated classes not shown

Figure S10

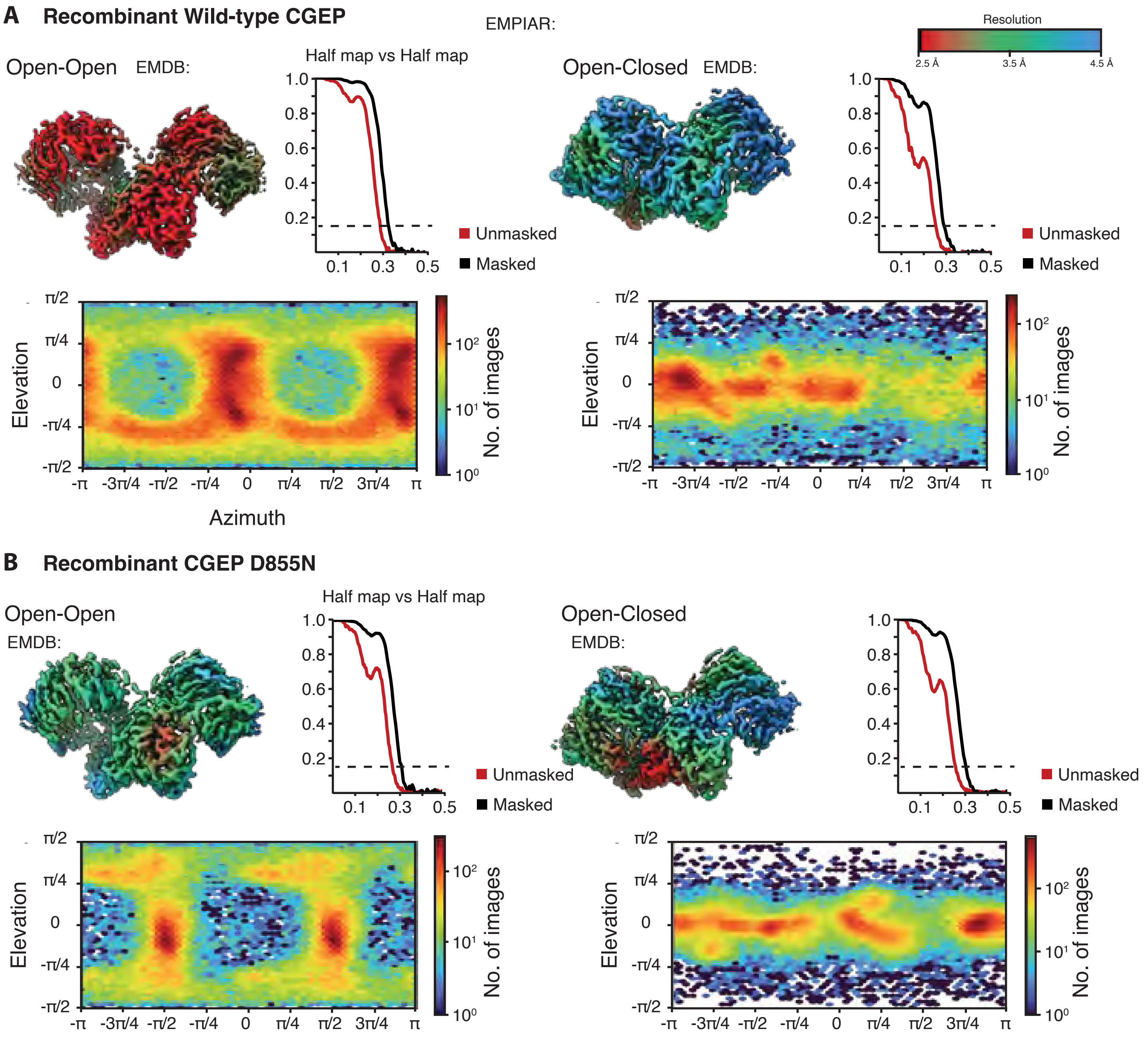

A
Recombinant Wild-type CGEP
EMPIAR:
Resolution
2.5 Å
3.5 Å
4.5 Å
Open-Open
EMDB:
Half map vs Half map
1.0
0.8
0.6
0.4
0.2
0.1
0.3
0.5
Unmasked
Masked
Open-Closed
EMDB:
Elevation
π/2
π/4
0
-π/4
-π/2
-π
-3π/4
-π/2
-π/4
0
π/4
π/2
3π/4
π
Azimuth
No. of images
$10^2$
$10^1$
$10^0$
B
Recombinant CGEP D855N
Open-Open
EMDB:
Half map vs Half map
Unmasked
Masked
Open-Closed
EMDB:
Elevation
No. of images

Figure S11

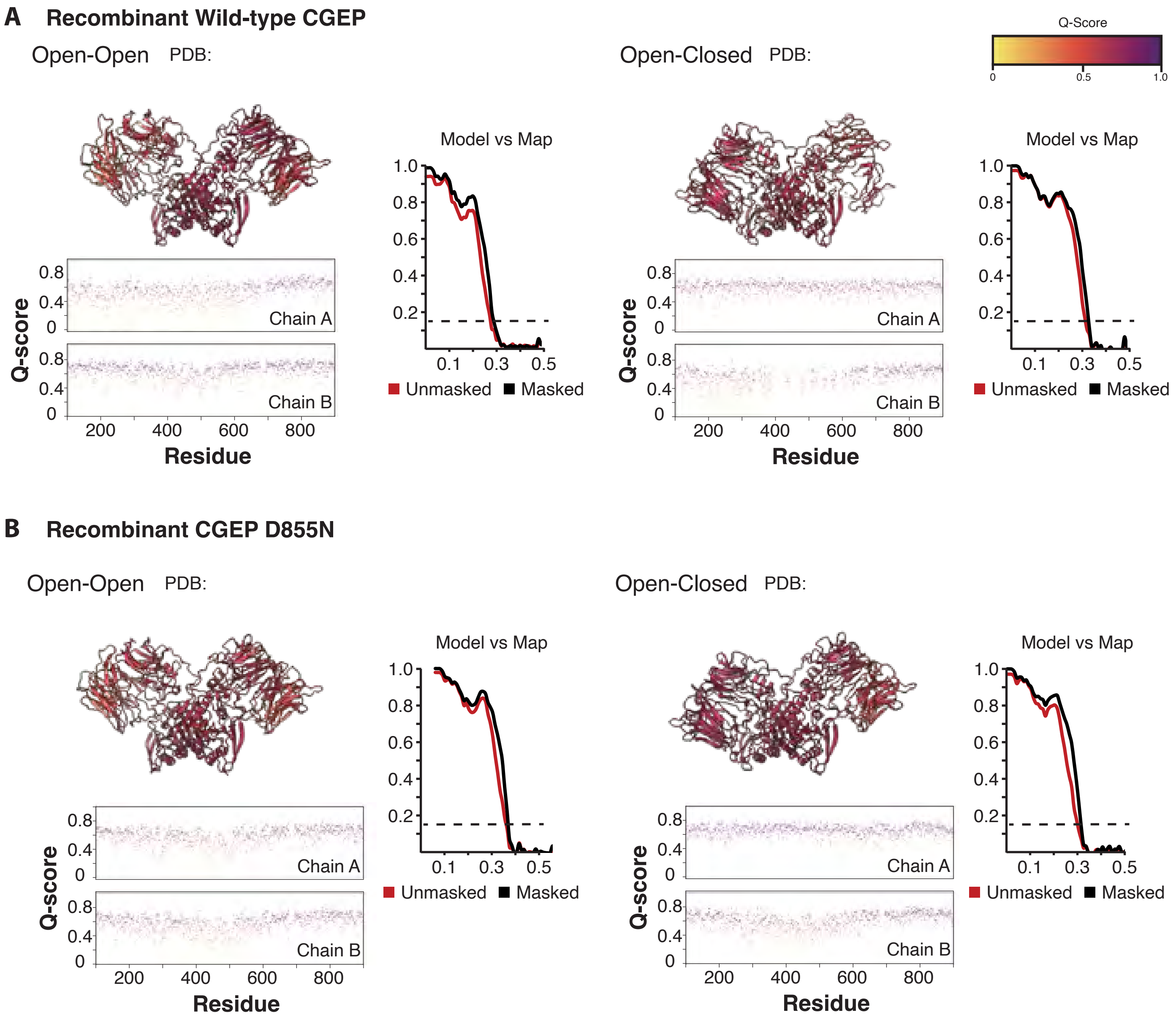
A Recombinant Wild-type CGEP
Open-Open PDB:
Open-Closed PDB:
Q-Score
0
0.5
1.0
Model vs Map
1.0
0.8
0.6
0.4
0.2
0.1
0.3
0.5
Unmasked
Masked
Q-score
0.8
0.4
0
Chain A
Chain B
200
400
600
800
Residue
B Recombinant CGEP D855N
Open-Open PDB:
Open-Closed PDB:
Model vs Map
Unmasked
Masked
Q-score
Chain A
Chain B
Residue
